# Prenatal alcohol exposure promotes nerve injury-induced pathological pain following morphine treatment via NLRP3-mediated peripheral and central proinflammatory immune actions

**DOI:** 10.1101/2025.05.30.655639

**Authors:** Andrea A. Pasmay, Ariana N. Pritha, Justin Carter, Alissa Jones, Annette K. Fernandez-Oropeza, Melody S. Sun, Diane C. Jimenez, Minerva Murphy, Carlos F. Valenzuela, Shahani Noor

## Abstract

Adverse in-utero conditions may exert a lifelong impact on neuroimmune function. Our prior work showed that prenatal alcohol exposure (PAE) increases pathological pain sensitivity (allodynia) following peripheral sciatic nerve injury. While the immune mechanism(s) of PAE-induced immune dysfunction are poorly understood, prior studies implicated the involvement of Toll-like receptor 4 (TLR4) and the nucleotide-binding domain, leucine-rich repeat-containing family, pyrin domain-containing 3 (NLRP3) inflammasomes. Interestingly, emerging data suggest a surprising overlap of spinal glial proinflammatory activation via the TLR4-NLRP3-interleukin (IL)-1β axis due to opioid treatment in nerve-injured non-PAE rodents. Considering this preclinical evidence, we explored whether PAE poses a risk factor in creating proinflammatory immune bias consequent to opioid (morphine) exposure. We hypothesized that under nerve injury conditions, PAE may interact with morphine, promoting peripheral and CNS proinflammatory factors in a NLRP3-dependent manner. Using a minor nerve injury model in adult mice, we demonstrate that PAE prolongs the chronicity of ongoing allodynia in both sexes, with a more pronounced effect observed in male mice. Our study shows that PAE amplifies proinflammatory responses at the injury site and the spinal cord, driving morphine-prolonged allodynia through NLRP3 inflammasome activation. Furthermore, high mobility group box 1 (HMGB1), a well-established pain-promoting TLR4 agonist, is elevated in allodynic PAE mice. NLRP3 inhibitor, MCC950, effectively reverses morphine-induced allodynia and reduces Caspase-1 activity, IL-1β, and related proinflammatory factors. Although few sex-specific effects were observed, our data convincingly support that PAE and morphine interactions ultimately converge on NLRP3-driven mechanisms in both sexes. Together, this study suggests that PAE modulates later-life neuroimmune function and provides critical insights into immune regulators underlying PAE-induced biological vulnerability to pathological pain processing and adverse effects of opioids.

## Introduction

Prenatal alcohol exposure (PAE) results in a range of adverse physiological and neurobehavioral outcomes, recognized as fetal alcohol spectrum disorders (FASDs) ^1,2^. FASDs occur worldwide and are as high as 5% in the US, estimated to be more frequent than other neurodevelopmental disorders, including autism ^2^. Notably, sensory processing issues ^3–5^, including touch hypersensitivity, are common in individuals with FASD. Consistent with clinical observations, numerous preclinical studies indicate that PAE induces biological vulnerability to lifelong dysfunctional central nervous system (CNS)-immune interactions ^2,6^ heightens proinflammatory neuroimmune activity upon exposure to subsequent immune stimuli ^7^, although key immune signaling pathways involved are poorly understood. In recent years, we have utilized an adult-onset peripheral nerve injury model to unmask PAE-induced susceptibility to chronic CNS dysfunction leading to pathological touch sensitivity (allodynia) ^4,8^.

Mechanical allodynia is a common clinical symptom of chronic pain. Chronic allodynia results from heightened excitation of spinal cord pain neurons due to aberrant neuronal-glial immune interactions ^9^, often consequent to peripheral nerve injury ^10^. Following nerve injury, endogenous cell stress factors (damage-associated molecular patterns, DAMPs) e.g., HMGB1, are released that, in turn, activate the immune receptor Toll-like receptor 4 (TLR4) on peripheral immune and spinal glial cells ^11–13^. TLR4 activation leads to translocation of the transcription factor, NF-kB, inducing the transcription of proinflammatory factors, including pro-IL-1β, pro-IL-18, TNF-α and nod-like receptor family pyrin domain containing 3 (NLRP3). NLRP3 further assembles with other proteins to form the NLRP3 inflammasome complex that activates Caspase-1, a necessary step for the production and release of a mature form of IL-1β ^14^ and IL-18 ^15–17^. Proinflammatory immune activation in the periphery results in the sensitization of nociceptors, resulting in increased excitability and peripheral sensitization ^18,19^. This enhanced nociceptive input is relayed to the spinal cord, where it activates glial cells and induces local inflammatory cascades, promoting central sensitization ^20,21^. Recent studies also highlight the involvement of the TLR4-NLRP3 axis in supraspinal regions, particularly in the midbrain periaqueductal gray (PAG) and the anterior cingulate cortex (ACC), which play critical roles in pain processing ^22–25^, reinforcing proinflammatory neuroimmune signaling along the pain pathway ^26–28^. Together, persistent TLR4-NLRP3-driven signaling cascade across the periphery, spinal cord, and brain contributes to the development and maintenance of mechanical allodynia ^1^.

Beyond their crucial neuroimmune modulatory roles during chronic allodynia, glia also plays a significant roles in influencing opioid tolerance and hyperalgesia associated with opioid pain therapeutics ^29–32^. While opioids, such as morphine, primarily act on the excitability of the pain neurons via mu-opioid receptors, inducing pain relief, opioid-induced proinflammatory actions via the TLR4-NLRP3-IL-1β pathway have been reported ^33^. In fact, proinflammatory effects from repeated morphine on primed immune cells ^34^ can act as an exogenous TLR4 activator underlying opioid-induced tolerance, and hyperalgesia observed clinically and in numerous preclinical pain models ^33,35,36^. While most preclinical studies examine morphine’s effects within hours of treatment, morphine paradoxically prolongs nerve injury-induced allodynia in male rats *after discontinuation* of treatment ^36,37^. While these studies highlighted the involvement of TLR4 and NLRP3 activation on spinal glia ^36^, potential sex differences and the effects of opioids modulating these immune factors in the periphery and brain regions remain unknown.

Our prior work suggests a potential overlap between neuroimmune pathways in morphine-mediated actions on glial cells ^38^ and PAE-induced changes in neuroimmune function ^6,39,40^ during adulthood. Notably, PAE rodents exhibit chronic allodynia following minor nerve injury, whereas non-PAE rodents subjected to the same injury do not ^39–42^. PAE-related susceptibility to chronic allodynia was associated with enhanced IL-1β and TNFα at the peripheral nerve injury site ^43^ and spinal cord ^41^. Our recent report explored the potential overlap of PAE-induced neuropathic pain susceptibility and morphine-mediated actions in female PAE mice ^44^. A striking observation was made suggesting that following minor nerve injury, the chronicity of allodynia is further exacerbated from a 5-day regimen of morphine treatment in PAE mice, whereas non-PAE control mice did not show allodynia. Moreover, utilizing a well-established small molecule inhibitor of NLRP3, MCC950 ^28,45–50^ we found that prolonged allodynia following morphine treatment was NLRP3-dependent. However, a comprehensive understanding of molecular changes and potential sex differences related to TLR4 and NLRP3 signaling driving this prolongation of allodynia under PAE conditions is yet to be explored.

Here, utilizing a previously characterized model of a minor sciatic nerve injury^1^, we hypothesized that PAE interacts with morphine to promote peripheral and CNS proinflammatory factors, driven by NLRP3 inflammasome activation. To test this hypothesis, we: **1)** Investigated whether PAE-induced susceptibility to morphine-prolonged allodynia occurs in both male and female mice; a side-by-side comparison characterizing the onset, duration and spontaneous reversal of allodynia was conducted. **2)** Performed behavioral assessments of allodynia, with or without MCC950-treated mice, to confirm the necessary role of NLRP3 in morphine-induced prolonged allodynia in both sexes. **3)** Examined molecular changes associated with the TLR4-NLRP3 axis in the pain-relevant PNS (sciatic nerve) and CNS (lumbar spinal cord and brain) regions.

## 2. Materials and Methods

### 2.1 Animals

All procedures were approved by the Institutional Animal Care and Use Committee (IACUC) of the University of New Mexico (UNM) Health Sciences Center. Animal experiments adhered to ARRIVE guidelines and the National Institutes of Health guide for laboratory animal care and use (NIH Publications No. 8023, revised 1978). All mice were routinely monitored by the animal care staff under the direction of the institutional veterinarian and staff, with cages and bedding changed every 7 days. All behavioral assessments, injections, and tissue collections were performed within the first 2 hours of the inactive cycle (light cycle) to avoid the influence of endogenous circadian regulation in the production of proinflammatory cytokines.

### 2.2 Prenatal alcohol exposure (PAE) paradigm

A well-established paradigm for generating experimental moderate prenatal alcohol-exposed offspring was utilized ^51,52^. PAE or age-matched control mice were provided at weaning by the New Mexico Alcohol Research Center (NMARC). Briefly, C57BL/6J mice were obtained from The Jackson Laboratory (Bar Harbor, ME) for breeding. Upon arrival, sires and dams were acclimated in a colony on a 12:12-hour reverse light/dark schedule and fed Teklad 2920X rodent chow and tap water, available *ad libitum*. On day 7, dams are isolated; on days 14-17, dams are exposed to 0.066% (w/v) saccharin (sac offspring, non-PAE control mice) or 5% w/v ethanol sweetened with 0.066% (w/v) sac solution (PAE offspring) for 4hr/day (1000 to 1400 hr). On day 18, dams are switched to 10% w/v ethanol sweetened with 0.066% (w/v) sac solution (PAE offspring). For breeding, individual females were placed into a cage with a single-housed male for two hours (from 1400 to 1600 hr) for three consecutive days. Pregnancy was positively determined by monitoring weight gain every 3-4 days. Alcohol exposure occurs throughout pregnancy, and this protocol produces blood EtOH concentrations of about 80 mg/dl following the drinking period in dams, with no negative effects on pup survival, weight, or litter size ^51^. One day after birth, access to alcohol was withdrawn using a step-down procedure, as described previously ^52^. saccharin (sac) (control mice) and PAE offspring were weaned at ∼3 weeks and subsequently maintained in groups of 2-4 mice per cage. Adult (7-8 months old) female & male PAE or age-matched sac (control) mice offspring were used for all experiments. None of the experimental groups contained more than two subjects from a given litter to avoid “litter effects.” Offspring were habituated to a standard light/dark cycle (lights on from 0600 h to 1800 h) for at least 3 weeks and kept in these conditions for the duration of the study.

### 2.3 Minor chronic constriction injury (CCI)

Mice were exposed to minor nerve injury (minor CCI) with a single suture (6–0 chromic gut) tied around the sciatic nerve, which is a modification of the previously well-established standard chronic constriction injury that involves more robust sciatic nerve damage, with three segments of 4-0 or 5-0 sutures ^44^. Following isoflurane anesthesia (induction at 1.5%-2.5 vol.% followed by 2.0 vol.% in oxygen), the dorsal left thigh was shaved and cleaned using 80% EtOH. The left sciatic nerve is gently exposed using blunt dissection scissors and isolated with sterile plastic probes. A single, 6–0 chromic gut suture (Ethicon, Cat #: 796G) is tied around the sciatic nerve. The suture is composed of a double knot located at the middle of the nerve without pinching the nerve to avoid nerve damage. The nerve was kept moist using isotonic sterile saline. Sham surgeries consist of an exposure of the nerve without any suture ligation. The nerve is carefully placed back into its position, and the overlying muscle is sutured using a 4–0 silk suture. The skin was closed using three Reflex™ wound clips (Kent Scientific Corp.; Cat #: INS750344). The total time for the surgical procedure was ∼15 min, followed by a ∼5-minute recovery from anesthesia. After the surgical procedure, the healing of the wound, hind paw autotomy, activity levels, and grooming appearance were assessed twice a day for three days to ensure a smooth and healthy recovery process. Staples were removed 10-12 days after surgery, at which point the wound had completely closed. No mice were excluded due to surgical complications.

### 2.4 Behavioral assessment of mechanical allodynia

Behavioral assessment of mechanical allodynia was performed using the Von Frey Fiber test, as described in our prior reports ^39,41,42^. Mice were gently handled and habituated to testers and the testing environment. Mice were tested in the same room as they are housed, limiting environmental variation. Mice were habituated to the testing environment for ∼60 min over 4 days before baseline assessment. The Von Frey test involves using nine calibrated monofilaments, which are applied for up to 3.0 seconds on the plantar surface of both the left and right hind paws. The testing of hind paw laterality is randomized, and the intertrial stimulus interval was at ∼10 seconds. Each paw undergoes a maximum of seven stimulus presentations. Lifting, jumping, licking, or shaking the paw was considered a positive response. The total number of positive and negative responses was then entered into the PsychoFit software to determine the absolute withdrawal threshold (50% paw withdrawal threshold). The PsychoFit program fits a Gaussian integral psychometric function to the observed withdrawal rates for each monofilament using a maximum-likelihood fitting method^53^. Cohorts of 6–8 mice were tested, with experimenters unaware of the experimental conditions. Each group consisted of at least 1–2 mice per treatment condition. Time points were carefully selected based on prior studies to capture allodynia’s development, persistence, and resolution ^40^. Behavioral evaluations were conducted at baseline (BL), post-surgery days 3, 7, 10, 14, and then every 2-4 days following morphine injection and 90 min and 1 day post MCC950 treatment.

### 2.5 Morphine treatment

Single subcutaneous (Sub-Q) injections of morphine were given for five consecutive days starting on D14 through D18 post-CCI. The timeline for morphine treatment was based on our previous characterization demonstrating that PAE mice reliably develop allodynia following minor nerve injury ^44^, with peak and stable mechanical allodynia occurring around D14. Injections were performed at the same time each day to ensure consistency and minimize variability. The analgesic isomer of morphine, (−) – morphine, was purchased from Sigma, MO, USA (Morphine sulfate salt pentahydrate, Cat #: M8777). Either morphine or vehicle (sterile saline) was injected at a dose of 10 mg/kg body weight, which is considered a moderate dose of morphine ^54,55^.

### 2.6 NLRP3 inhibitor, MCC950, treatment

MCC950, a small molecule inhibitor, was used to assess the role of NLRP3 in the development of morphine-prolonged allodynia. MCC950 inhibits NLRP3 oligomerization by preventing the assembly of ASC (apoptosis-associated speck-like protein) to the NLRP3 complex ^45^ and the release of mature IL-1β^56^. To date, numerous *in vivo* and *in vitro* studies have validated that MCC950 crosses the blood-brain barrier ^6^ and specifically blocks the NLRP3 inflammasome activation, not other inflammasomes that may induce Caspase-1 ^57^. Moreover, MCC950 has been successfully used to reverse chronic allodynia in numerous preclinical models, without affecting normal nociception ^58–61^. Using aseptic procedures, a single intraperitoneal injection of MCC950 (10 mg/kg) or vehicle injection was given to unanesthetized mice. The MCC950 dose was chosen based on prior in vivo studies demonstrating reduced neuroinflammation ^47,56,62^. The total time required for handling and injection procedure was less than 1 minute.

### 2.7 Tissue collection for RNA and protein analysis

Immediately after the last behavioral assessments (D24), tissues were collected for molecular analysis. The mice were induced into deep anesthesia using isoflurane (10 minutes at 5 vol.% isoflurane in oxygen at a 2.0 vol.%, then underwent rapid transcardial perfusion with ice-cold 0.1 M phosphate-buffered saline (PBS; pH = 7.4; flow rate 10 mL/min). Blunt dissection scissors were used to re-expose the injured sciatic nerve (SCN, left side). Approximately 1 cm of the SCN was collected, keeping the suture (lesion site) in the middle. Spinal cord was gently flushed out from the vertebral column with ice-cold sterile PBS into a petri dish and ipsilateral lumbar spinal cord dorsal horn LSC (L3-L6) was dissected. The brain was carefully removed from the skull. Midbrain was dissected from the entire brain and the anterior cingulate cortex was dissected from the contralateral (right) hemisphere. All tissues were immediately flash-frozen in dry ice in DNase/RNase/Protease-free 1.5 mL Eppendorf tubes (VWR International; Cat #: 20170-038) and stored at −80 °C for future analysis.

### 2.8 Total RNA & protein isolation

Tissues were processed for RNA and protein analysis, as described previously ^42,43^. Tissues were homogenized with sterile RNAase-free PBS using a motorized VWR disposable pellet mixer and cordless motor pestle system. Homogenized tissue was divided by volume to a ratio of 60:40 and used for RNA and protein analysis, respectively. RNA extraction was performed using Qiazol Lysis Reagent and the miRNeasy Micro Kit (Qiagen; Cat #: 74004), per the manufacturer’s instructions. The concentrations and quality of the total RNA were assessed by NanoDrop (Thermo Scientific, MA, USA). Tissue homogenates designated for protein analysis were preserved in a protein buffer containing protease inhibitor (VWR; Cat #: 7844) and lysis buffer (Thermo Scientific; Cat #: P187787). The tissue samples were homogenized, sonicated, and centrifuged at 14,000g for 10 mins to separate pellet debris from the protein lysate. Per the manufacturer’s instructions, the total protein concentration was determined using a Bradford Assay (Bio-Rad Cat #: 5000201).

### 2.9 High-sensitivity IL-1β ELISA protein Assay

Protein concentration was determined using the Quick Start Bradford Protein Assay (Bio-Rad, USA, Cat #: 5000201). IL-1β protein was detected using a high-sensitivity commercial enzyme-linked immunosorbent assay kit (HS Quantikine; R&D Systems, USA, Cat #: MHSLB00) according to the manufacturer’s instructions. IL-1β concentrations were calculated against the internal standards and normalized to the total protein concentration in the homogenate (pg/mL). Due to low protein yield achieved from designating each tissue sample for both protein and mRNA analysis, some samples were pooled from multiple biological replicates and few samples were only designated for protein analysis. For the spinal cord IL-1β assay, a total of 100 µg (females) or 65 µg (males) protein was used. For the sciatic nerve IL-1β 90 µg of total protein was used in both males and females.

### 2.10 Caspase-1 activity assay

Caspase-1 activity is an immediate downstream effect of NLRP3 inflammasome activation. Colorimetric detection of Caspase-1 activity assay was performed according to the manufacturer’s instructions (Abcam Cat #: ab273268). Most quantitative molecular assays detect both pro-IL-1β and mature forms of IL-1β protein from tissue lysates. Caspase-1 activity assay utilizes the activity of Caspase-1 that recognizes the sequence YVAD, hence providing a reliable readout of NLRP3 inflammasome activation and production of mature IL-1β ^63^. Briefly, protein samples were measured by Bradford Assay, and 60-65 µg total protein was incubated with 5 µl of YVAD-p-nitroanilide (pNA) and incubated at 37°C for 2 hours. The activity of Caspase-1 is measured by spectrophotometric detection of the chromophore p-nitroanilide (pNA) at 400 nm after cleavage from the labeled substrate YVAD-pNA. For spinal cord tissue, all experimental groups were included for females, but due to the limited protein amount left from male tissues, additional mice were added to the study, with only the three critical groups.

### 2.11 mRNA analysis by Quantitative Real-Time PCR

Relative mRNA levels were measured as described in our prior reports ^42^. Briefly, total RNA samples were diluted to a standardized RNA concentration, and their concentrations and quality were further confirmed by NanoDrop measurement. Total RNA (1.0-1.2 μg) was used to synthesize reverse transcription (cDNA). For cDNA synthesis, SuperScript™ IV VILO™ cDNA Synthesis Kit (Thermo Fischer Scientific, Cat #: 11754050) was used per the manufacturer’s instructions. A 1:2.5 dilution factors were applied to cDNA samples for assessment of transcripts of interest in the ipsilateral spinal cord, ipsilateral sciatic nerve, contralateral ACC, and whole midbrain. The 1:200 dilutions of cDNA were used to assess the normalizer transcripts (18s RNA) for each tissue sample. All samples were assayed in triplicates via quantitative real-time PCR (qRT-PCR) with TaqMan Gene Expression Assays^42^. Standard deviations were calculated for triplicate measurements, and when exceeding 0.1, the average value of the two closest replicates was utilized. Relative gene expression levels were analyzed based on the endogenous controls (normalizer, 18s), and data are presented as fold increases relative to the non-allodynic control group (sac +minor CCI+ veh + veh), using the formula: 2^-ΔΔ*CT*^ ^64^. The following pain-relevant proinflammatory transcripts were evaluated: high mobility group box 1 (HMGB1, hmgb1) interleukin-1β (IL-1β, *il1b*), interleukin-18 (IL-18, *il18*) tumor necrosis factor α (TNFα, *Tnf*), NOD-like receptor protein-3 (NLRP3, *nlrp3*), IkappaB alpha (IKBA. *lkBa*), GFAP (*Gfap*), an astrocyte activation marker, and Iba-1 (*Aif-1*), a marker for microglia activation. The μ-opioid receptor is known to play a crucial role in modulating pain signaling pathways ^65^. Although it is beyond the scope of this study to examine activation of the u-opioid receptor from morphine, expression of the μ-opioid receptors was measured to examine potential PAE-related effects. Samples from male and female mice were processed simultaneously to minimize variability.

### 2.12 Experimental design

Based on the clinical relevance and strong scientific premise from prior work, we focused on mechanical allodynia in this study, while other pain modalities such as mechanical or thermal hyperalgesia, may also be influenced via PAE and morphine interaction and differentially regulated by MCC950 ^28^, which warrants further investigation. Moreover, to avoid confounding factors from multiple behavioral assessments, we have not assessed these mice for other behavioral effects such as anxiety, locomotor activities ^66,67^ that may be influenced by PAE and morphine. A total of 153 adult mice, including sac (control) and PAE offspring, were used in hind paw sensitivity assessments reported in this study. To explore the effect of morphine during minor nerve injury, adult mice with prenatal alcohol exposure (PAE) and age-matched non-PAE (sac) mice underwent sham injury or minor CCI and received morphine or vehicle treatment. Hind paw sensitivity was assessed at various time intervals to track the progression of allodynia until mice exhibited hind paw sensitivity comparable to baseline levels (Figure 1). For Figure 1, a total of 58 mice, 29 male, and 29 female mice, were used, with an N= 4-5 per treatment group. To specifically assess the role of NLRP3 in the maintenance of morphine-prolonged allodynia beyond the period of spontaneous recovery (typically resolved by day 21 in PAE mice, shown by our previous work ^39^) MCC950 or vehicle treatment were administered on day 23, when persistent allodynia could be attributed to neuroimmune dysregulation rather than nerve injury alone. (Figure 2). For Figure 2, a total of 95 mice, 46 male, and 49 female mice, were used, 6-11 mice per treatment group. Based on our previous molecular comparisons (effect size, *f* = 0.5) ^39^, an a priori power analysis (G*Power, α = 0.05, power = 0.80, 8 groups) indicated a total sample size of 34 mice (∼6/group) to detect differences between allodynic and non-allodynic mice. Tissues were collected from all 95 mice, and molecular changes were assessed in each sample. Data are presented in Figure 3-9 and all supplemental figures. Two samples were excluded from the molecular analysis based on tissue collection/tissue processing notes suggesting potential tissue compromise.

**Figure 1.**
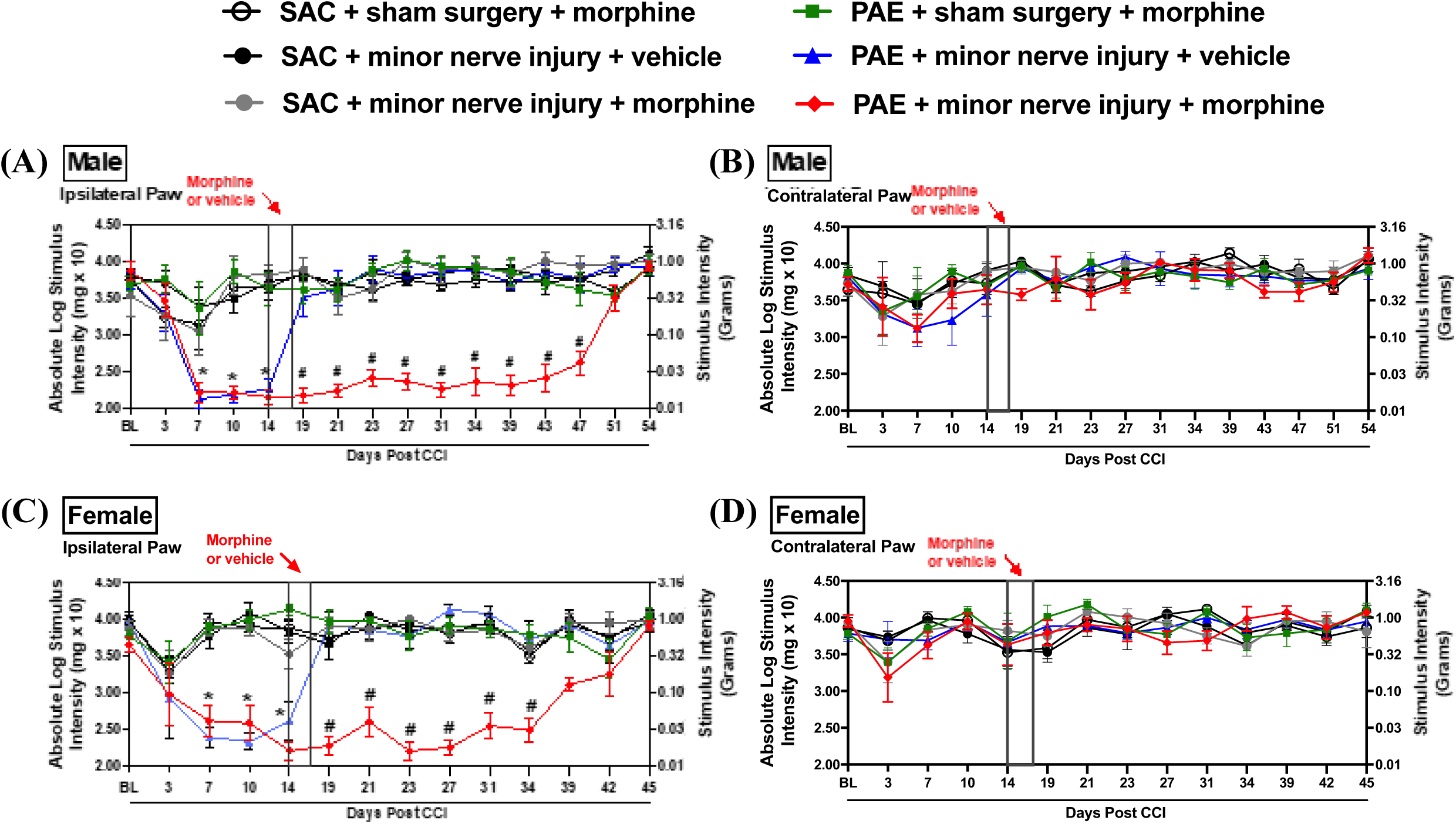
Morphine treatment prolongs the duration of allodynia only in minor nerve-injured PAE mice in both sexes. Sac and PAE mice of both sexes were exposed to sham surgery or minor nerve injury (CCI), and the effects of morphine treatment on hind paw sensitivity were examined. Pre-surgery baseline (BL) values were similar in sac and PAE mice (F_1,54_ = 0.48, p = 0.49). Ipsilateral hind paw sensitivity developed following minor CCI. **(A & C)** A significant interaction between PAE × time (F_3,150_ = 7.06, p = 0.00002), PAE × surgery (F_1,50_ = 104.71, p < 0.0001), and PAE × surgery × time (F_3,150_ = 4.03, p = 0.009) was observed only on the ipsilateral side. In males and females, nerve-injured PAE mice displayed a significant increase in ipsilateral hind paw sensitivity by D7–D14 (p < 0.05). **(B & D)** In both sexes, no contralateral allodynia was observed post-minor nerve injury. Post-morphine hind paw responses were evaluated between the PAE morphine-treated and vehicle-treated mice. **(A)** Post-morphine treatment, there was a significant interaction of PAE × morphine (F_1,29_ = 182.39, p < 0.0001). In males, morphine-injected PAE mice showed persistent ipsilateral hind paw sensitivity compared to vehicle-treated PAE mice up to D47 post-CCI (D23–D47; #p < 0.04). **(C)** Female PAE mice showed persistent ipsilateral hind paw sensitivity compared to vehicle-treated PAE mice up to D39 post-CCI (D19–D39; #p < 0.0001). **(A & C)** Morphine-injected PAE mice exhibited allodynia, in contrast to morphine-treated sac mice, which maintained baseline levels throughout the entire time course. **(B & D)** No significant changes were observed in the contralateral paw. Large effect sizes were observed for PAE exposure (η²ₚ = 0.870), morphine treatment (η²ₚ = 0.869), and their interaction (PAE × morphine; η²ₚ = 0.875), indicating that PAE mice treated with morphine developed significant allodynia compared to both sac morphine-treated and PAE vehicle-treated mice.

**Figure 2.**
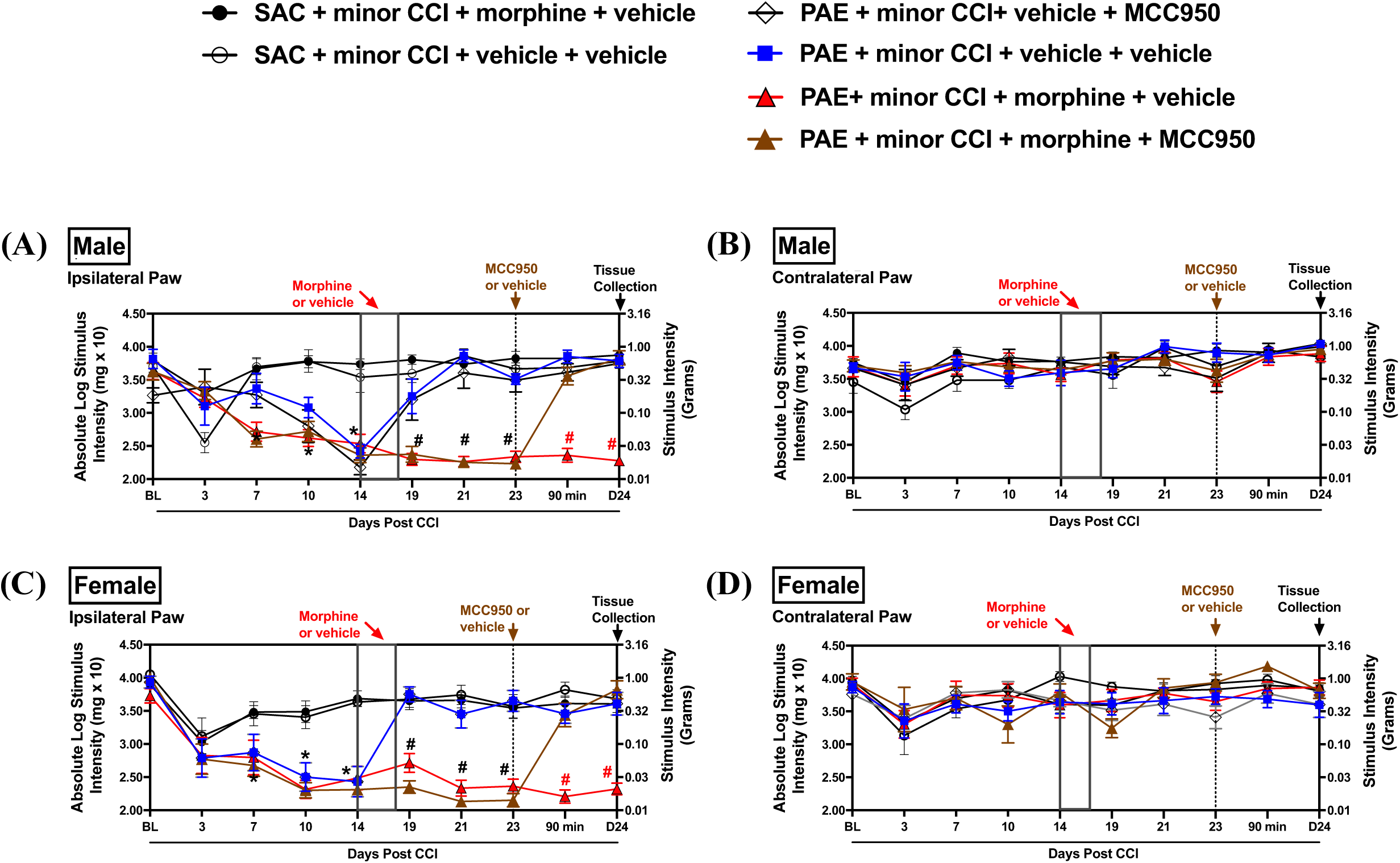
Systemic treatment of NLRP3 inhibitor, MCC950, reverses morphine-mediated prolonged allodynia in PAE mice, regardless of sex. The effects of systemic MCC950 treatment were examined during morphine-prolonged allodynia in nerve-injured PAE mice. A two-way ANOVA revealed no significant differences in pre-surgery (BL) values (F_1,91_ = 2.9, p = 0.09). Following CCI, only PAE mice developed morphine-prolonged allodynia, as indicated by a significant PAE interaction (F_1,364_ = 11.17, p = 0.0009). **Male (A)** and **female (C)** PAE mice developed allodynia in comparison to sac mice post-CCI at D7–D14 (p < 0.003). **(A & C)** Post-morphine treatment, there was a significant interaction of PAE × morphine (F_1,86_ = 132.77, p < 0.0001), as well as a sex × time interaction (F_2,172_ = 4.17, p = 0.0170). Morphine-treated PAE mice exhibited prolonged allodynia on D19–D23 (#p < 0.0001) compared to PAE vehicle-treated mice for both males and females. Additionally, regardless of sex, significant differences in allodynia were observed between morphine-treated PAE mice and morphine-treated sac mice at D19–D23 (p < 0.04). **(B & D)** Morphine treatment did not result in any changes in the contralateral hind paw. **(A & C)** There was a significant sex effect (F_1,60_ = 240.3, p < 0.0001) when evaluating the effects of MCC950. In both males and females, MCC950 treatment decreased sensitivity in morphine-treated PAE mice compared to vehicle-treated morphine-treated PAE mice, beginning as early as 90 minutes post-injection, with reversal continuing 24 hours post-injection (#p < 0.0001), at which time tissues were collected. **(B & D)** Regardless of sex, MCC950 did not alter hind paw responses in the absence of injury on the contralateral side.

**Figure 3.**
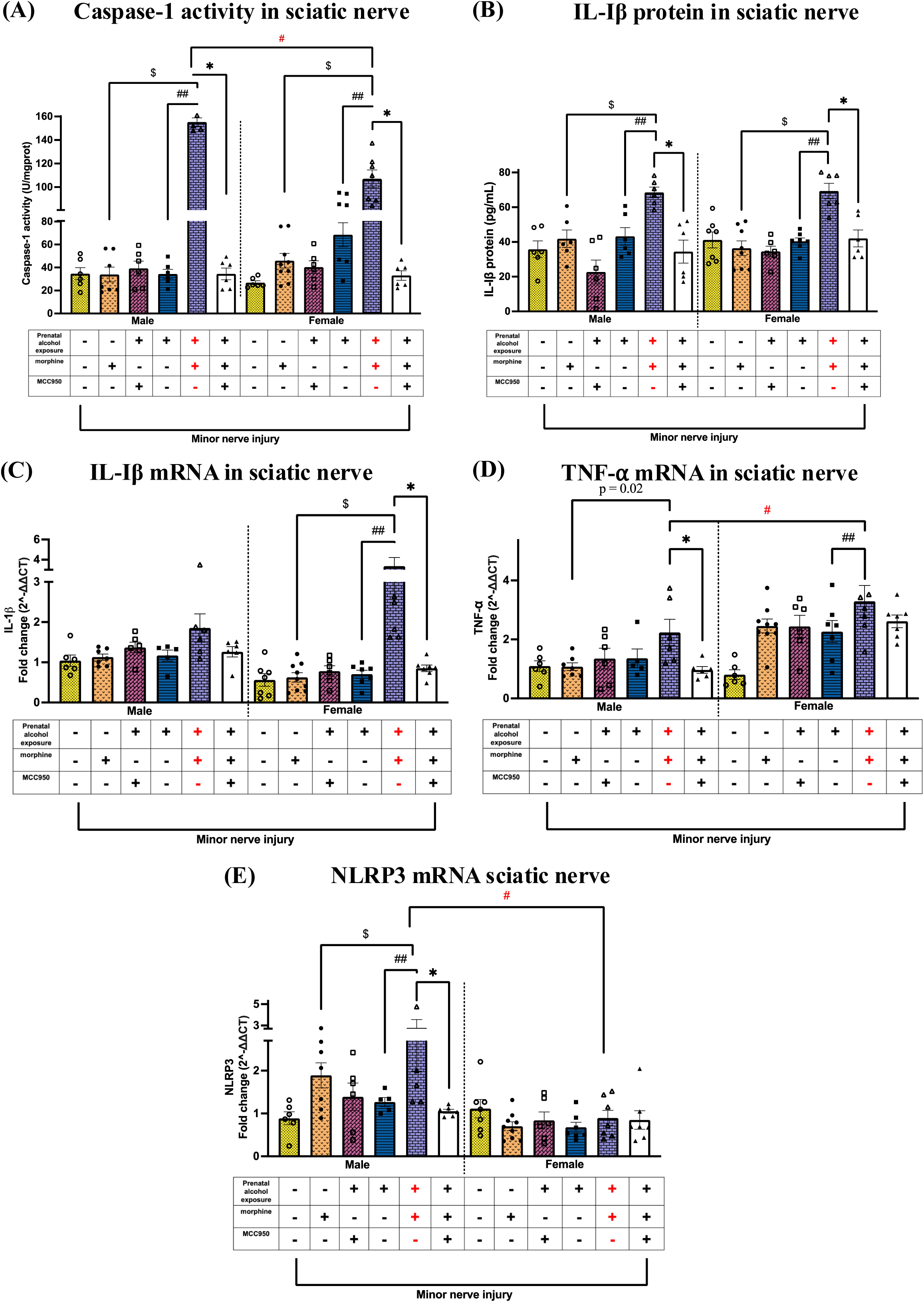
Effects of PAE and morphine immune interactions, and MCC950 treatment on Caspase-1, IL-1β, NLRP3, and TNF-α levels at the injured nerve. Ipsilateral sciatic nerves were collected from behaviorally verified mice, as represented in Figure 2. Prenatal exposure (+), morphine (+), and MCC950 (−) in purple denote the allodynic group; all other groups are non-allodynic or allodynia-reversed at this post-CCI time point. **(A)** Despite all mice being exposed to minor nerve injury, morphine treatment significantly increased caspase-1 activity regardless of sex in PAE mice compared to the two key control groups: sac male mice ($p < 0.0001, PAE vs. sac, morphine-treated mice) and PAE mice without morphine treatment (##p < 0.0001, PAE, morphine vs. vehicle). Post hoc comparisons revealed that male PAE morphine-vehicle mice exhibited significantly higher values compared to their female counterparts (#p < 0.0001). In both males and females, MCC950 treatment reduced caspase-1 activity in morphine-treated PAE mice that displayed allodynia reversal (*p < 0.0001). **(B)** Despite all mice being exposed to minor nerve injury, morphine treatment significantly increased IL-1β protein regardless of sex in PAE mice compared to the two key control groups: sac male mice ($p < 0.0009, PAE vs. sac, morphine-treated mice) and PAE mice without morphine treatment (##p < 0.004, PAE, morphine vs. vehicle). In males, MCC950 treatment reduced IL-1β protein in morphine-treated PAE mice that displayed allodynia reversal (*p < 0.04). **(C)** In females, morphine-prolonged allodynia coincided with a significant increase in IL-1β mRNA in PAE females compared to sac female mice ($p < 0.0001, PAE vs. sac, morphine-treated mice) and PAE mice without morphine treatment (##p < 0.0001, PAE, morphine vs. vehicle). MCC950 treatment reduced IL-1β mRNA in morphine-treated PAE females that displayed allodynia reversal (*p = 0.01). **(D)** In males, morphine-prolonged allodynia resulted in a significant increase in TNF-α mRNA levels exclusively in PAE mice treated with morphine compared to sac mice treated with morphine (p = 0.003, unpaired t-test). Morphine-prolonged allodynia resulted in a significant increase in TNF-α mRNA levels in PAE female mice treated with morphine compared to PAE females treated with vehicle (##p = 0.04, PAE, morphine vs. vehicle). MCC950 treatment reduced TNF-α mRNA in morphine-treated male PAE mice that displayed allodynia reversal (*p = 0.03). We observed significant sex differences in TNF-α mRNA levels, with females exhibiting higher levels than males in allodynic, morphine-treated PAE mice (#p = 0.04). **(E)** In male mice, morphine treatment significantly increased NLRP3 mRNA in PAE mice compared to the two key control groups: sac male mice ($p = 0.05, PAE vs. sac, morphine-treated mice) and PAE mice without morphine treatment (##p = 0.002, PAE, morphine vs. vehicle). NLRP3 mRNA was significantly higher in allodynic male morphine-treated PAE mice compared to their female counterparts (#p < 0.0001). MCC950 significantly reduced NLRP3 mRNA only in morphine-treated male PAE mice that displayed allodynia reversal (*p = 0.007).

### 2.13 Statistical analysis

#### Behavioral Data Analysis

All behavioral data were analyzed using GraphPad Prism (GraphPad Software Inc.; RRID: SCR_002798) or RStudio. For Figures 1 and 2, baseline comparisons were performed by collapsing groups across surgery, morphine, and MCC950 conditions, and a two-way ANOVA (sex × PAE) was used to assess group differences. To evaluate the development of allodynia after surgery, groups were collapsed by surgery condition (minor nerve injury vs. sham), and a three-way repeated measures (RM) ANOVA (sex × PAE × surgery × time) was conducted. Tukey’s post hoc tests were used to assess differences between PAE + minor nerve injury, sac + minor nerve injury, and PAE + sham groups. For Figure 2A, sham mice were not part of the design; therefore, a two-way RM ANOVA (sex × PAE × time) was used. Post-morphine time points were analyzed separately using a three-way RM ANOVA (sex × PAE × morphine × time), with Tukey’s test applied to assess whether PAE + minor nerve injury + morphine mice exhibited greater allodynia compared to sac + minor nerve injury + morphine and PAE + minor nerve injury + vehicle mice. Sham groups were excluded from this analysis. Potential sex differences in morphine response were also evaluated. The effects of MCC950 were analyzed using a two-way RM ANOVA (sex × MCC950 × time), with Tukey’s test applied to compare PAE + minor nerve injury + morphine + vehicle and PAE + minor nerve injury + morphine + MCC950 groups.

#### Molecular data analysis

Normality was assessed using the Shapiro-Wilk test. To maintain consistency across behavioral and molecular analyses, the same group variables and post hoc strategies were applied unless otherwise specified in the figure legends. Molecular outcomes related to morphine-prolonged allodynia were analyzed using a three-way ANOVA (sex × PAE × morphine), with Fisher’s LSD test used for post hoc comparisons. Sex differences in molecular responses were also evaluated. Molecular outcomes related to MCC950 effects were analyzed using a two-way ANOVA (sex × MCC950), with Fisher’s LSD post hoc tests. Adjusted *p*-values are reported, and all data are presented as mean ± SEM. In cases where the respective ANOVA did not yield significant values on post hoc comparisons from primary groups of interest unpaired t-tests was used based on our *a priori* hypothesis. Outliers were identified using Grubbs’ Test (GraphPad QuickCalc Outlier Calculator; α = 0.05). In figures depicting molecular data, Y-axes were broken to improve visualization across groups with widely varying expression levels. Main effects are reported in Table 1.

**Table 1:**
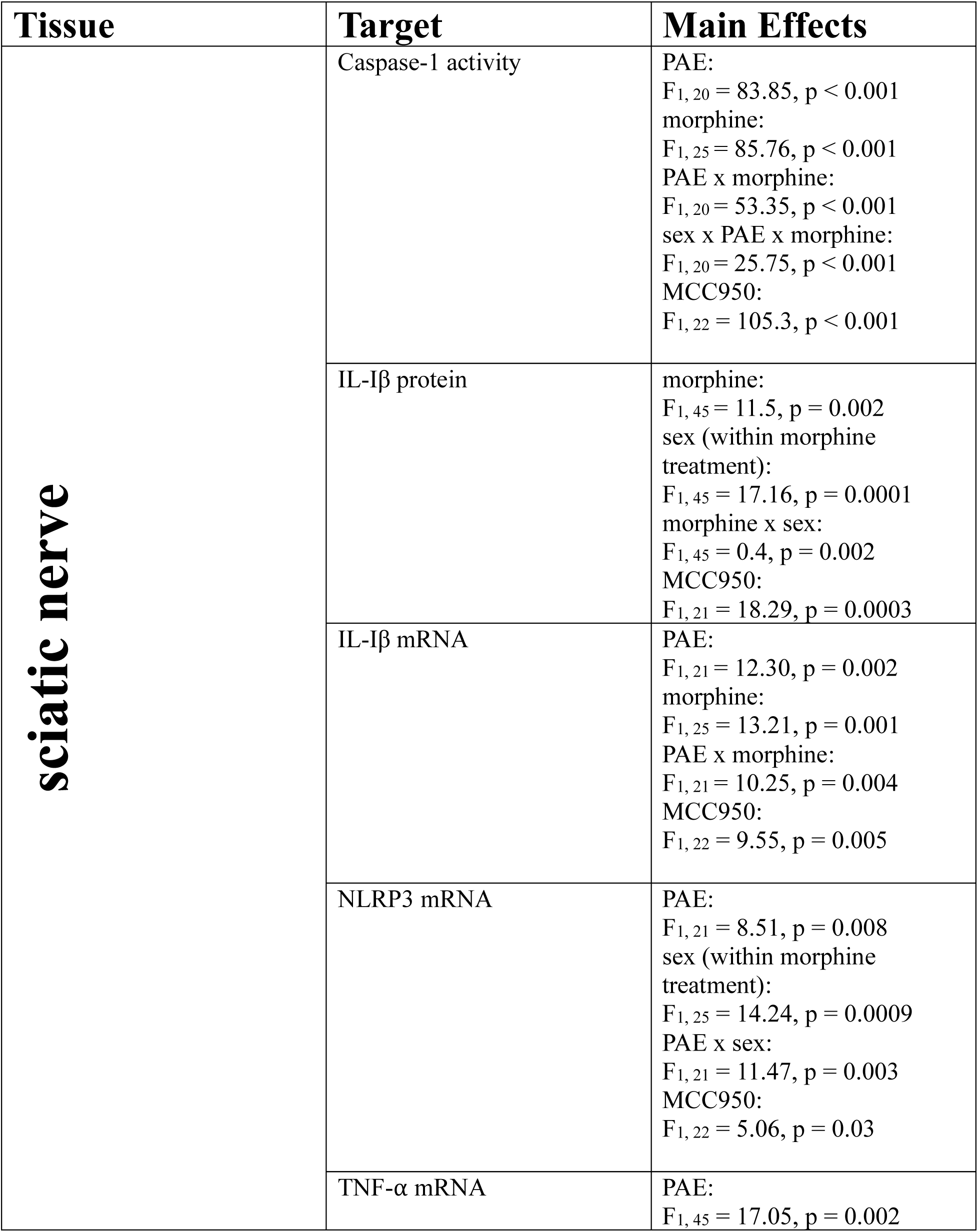
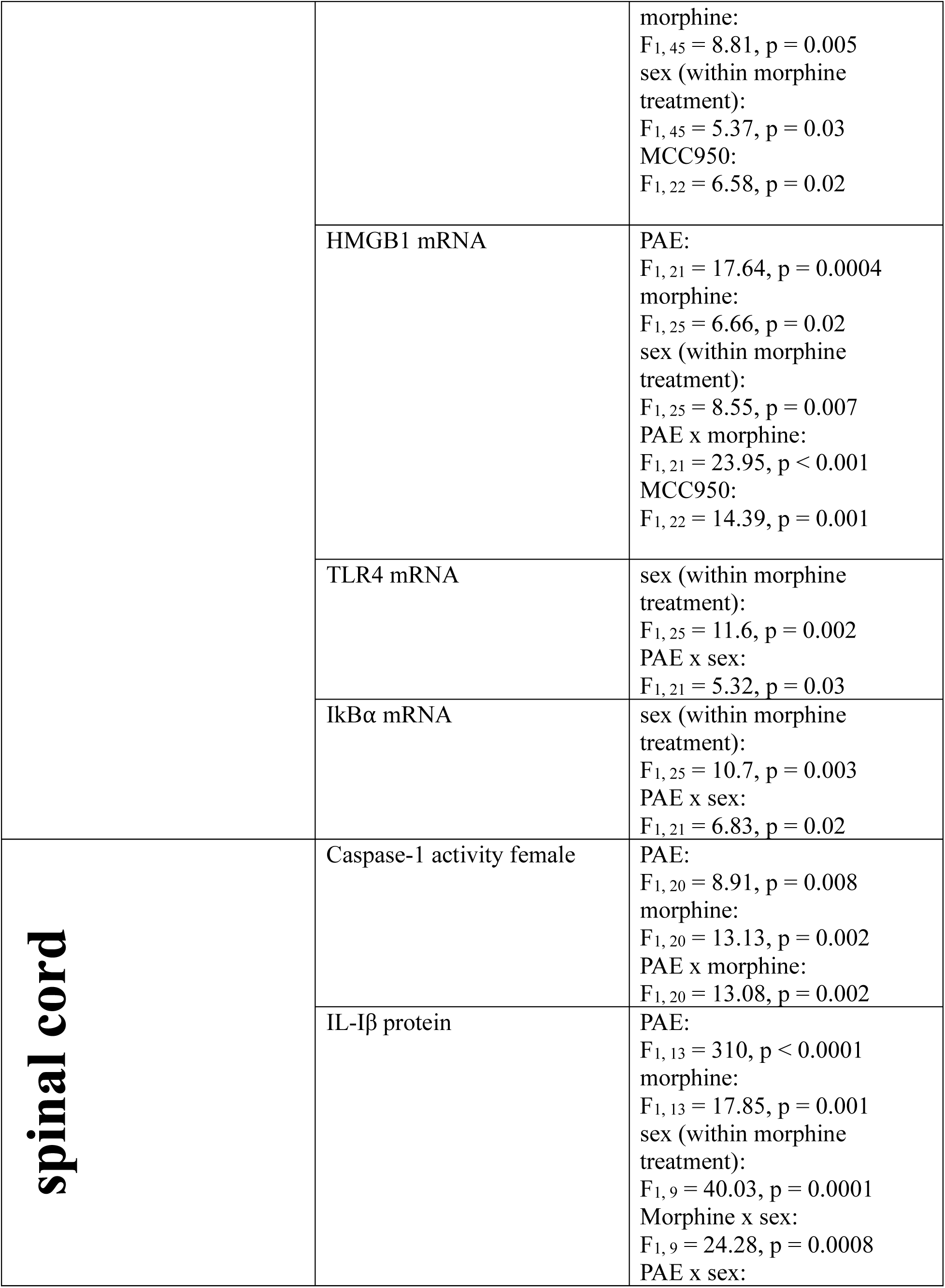
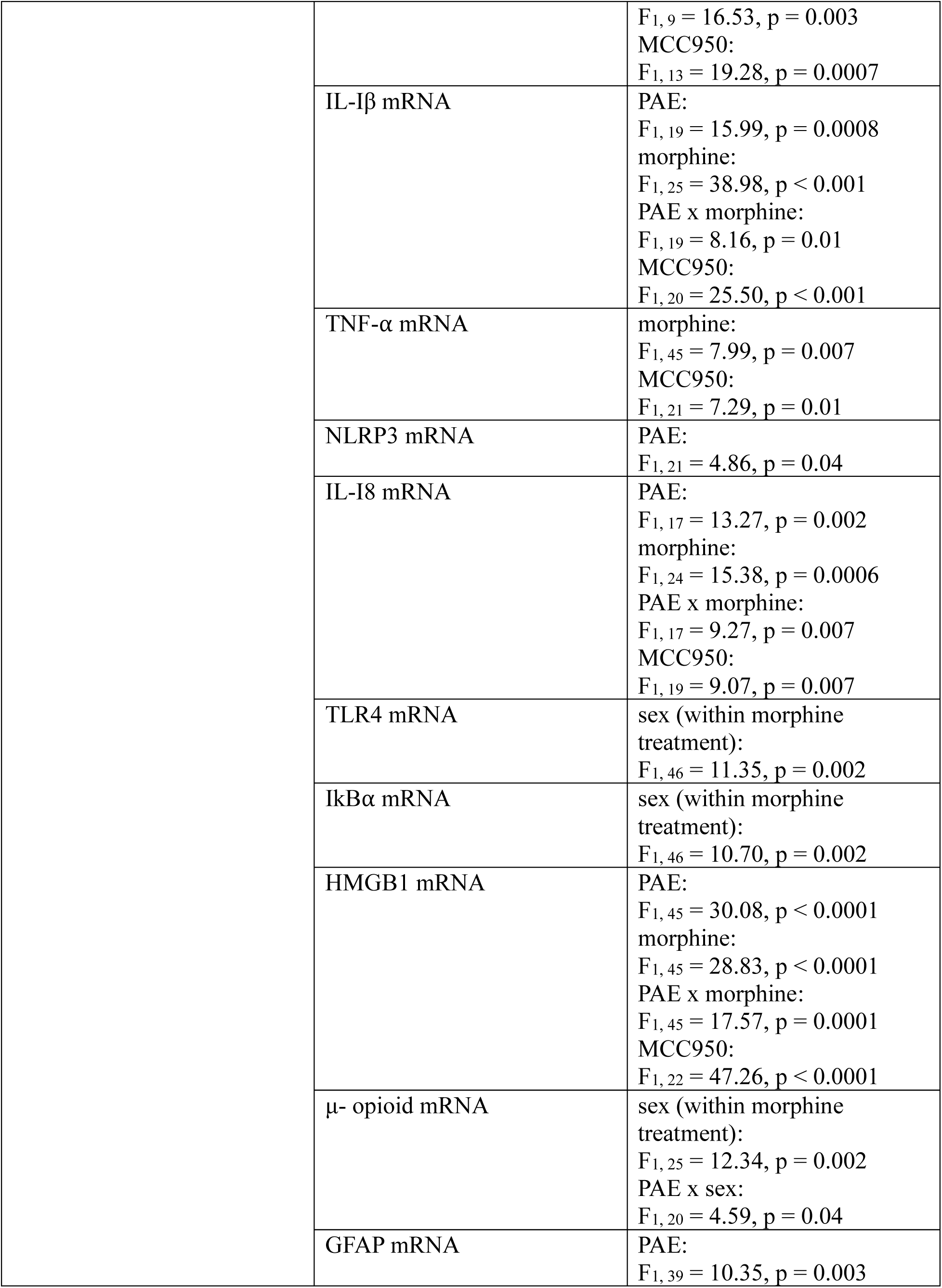
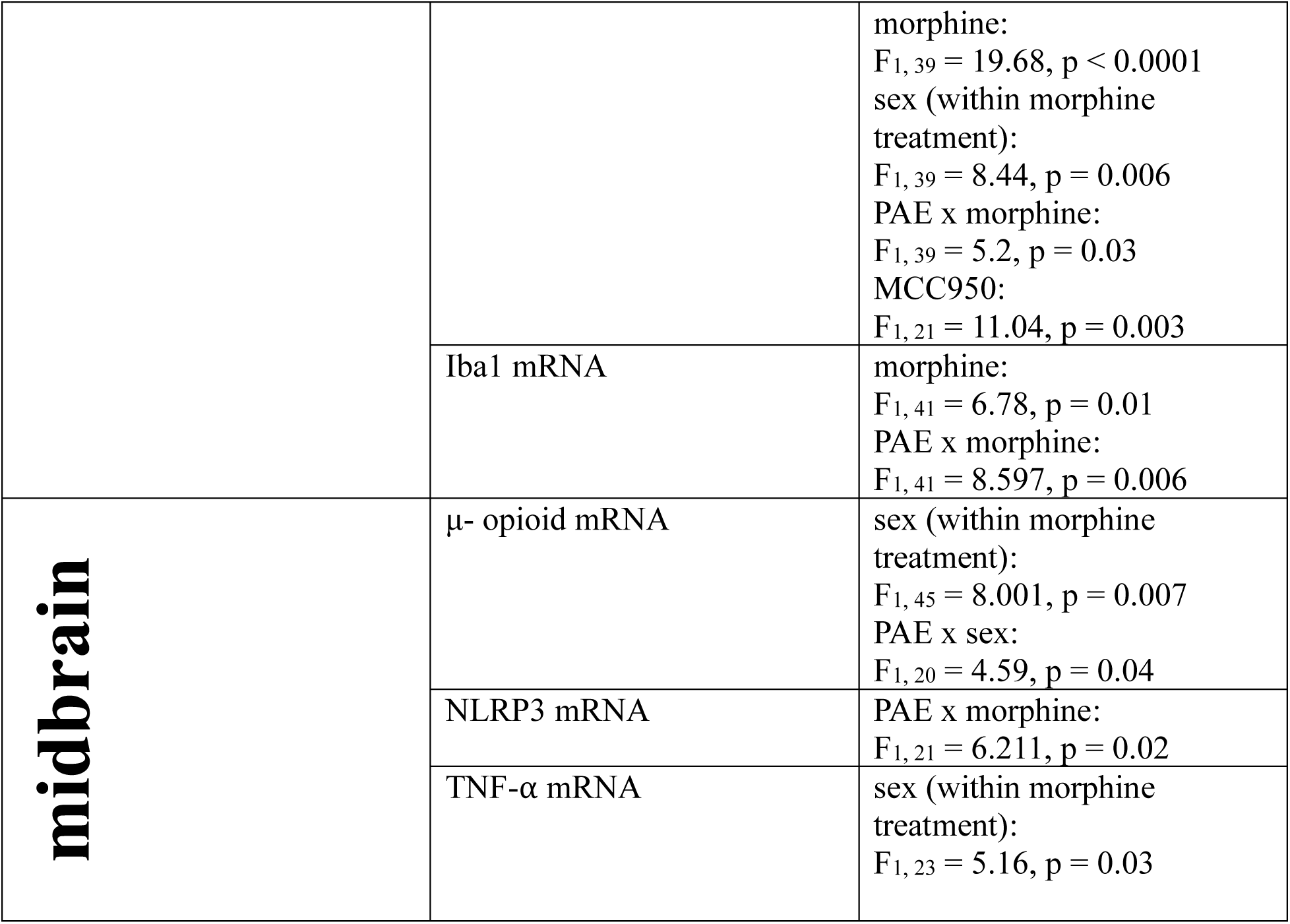
Key Molecular Findings: Main Effects.

## 3. Results

### 3.1 Repeated treatment of moderate doses of morphine results in the prolongation of minor nerve injury-induced allodynia, only in PAE mice

A well-established model of minor chronic constriction injury was utilized to examine the potential interaction of PAE and morphine. Adult PAE mice and non-PAE control (sac) mice were exposed to minor CCI or sham surgery. Prior to surgery, PAE and sac control mice displayed similar levels of baseline (BL) hind paw sensitivity. Following sham surgery, their hind paw sensitivities remained similar to pre-surgery BL values (**Figure 1**). Replicating our prior reports ^40^, minor CCI resulted in chronic unilateral allodynia only in PAE mice, whereas minor nerve-injured sac mice did not develop allodynia. A main effect of PAE (F₁,₃₀ = 33.54, p < 0.0001) was observed on hind paw responses ipsilateral to the sciatic damaged side from D1–D14 post-CCI time points; no sensitivity was detected in the contralateral side (**Figure 1A–B**). At D14 post-CCI, mice were treated with a moderate dose of morphine for five consecutive days. Mice were tested every 2–4 days until all mice displayed ipsilateral hind paw sensitivities similar to their BL values. Comparing hind paw responses at post-morphine time points in nerve-injured mice, a main effect of PAE (F₁,₃₀ = 200.75, p < 0.0001) was observed. In both male and female sac mice, morphine treatment did not increase ipsilateral or contralateral paw sensitivity. In contrast, minor nerve-injured male PAE mice continued to display unilateral allodynia until D47 for males and D34 for females post-CCI following morphine treatment, whereas vehicle-treated nerve-injured PAE mice spontaneously reversed by D19 (**Figure 1A, C**). A significant interaction effect of PAE × morphine (F₁,₂₉ = 182.39, p < 0.0001) was observed, with notable differences between allodynic morphine-treated PAE mice and vehicle-treated PAE mice from D19 to D34 in females and from D19 to D47 in males. In both male and female mice, no changes in hind paw sensitivity were observed in the contralateral hind paw at post-morphine treatment time points; nerve-injured PAE mice remained at about BL through the entire time course (**Figure 1B, D**). There was a significant sex × time interaction (F₆,₁₈₀ = 3.39, p = 0.0034), suggesting that males and females exhibit different trajectories of morphine allodynia development over time. A significant sex × PAE × time interaction was also observed (F₆,₁₈₀ = 2.90, p = 0.0102).

### 3.2 NLRP3 inflammasome is necessary for the development of morphine-induced prolonged allodynia in nerve-injured PAE mice

Utilizing the small molecule inhibitor MCC950, we explored the key role of NLRP3 inflammasome activation in driving morphine-induced allodynia under PAE conditions. Following the same experimental paradigm described in Figure 1, minor nerve-injured sac or PAE mice underwent morphine or vehicle treatment, followed by MCC950 administration. Reproducing the observations in Figure 1, both female and male minor nerve-injured PAE mice developed robust allodynia, whereas sac + minor CCI mice remained stably non-allodynic. A main effect of PAE was observed only on the ipsilateral side of the injury (F₁,₃₆₄ = 11.17, p < 0.0009; **Figure 2A, C**). In both sexes, morphine treatment further prolonged ongoing allodynia only in nerve-injured PAE mice (**Figure 2A, C**). Analyzing post-morphine treatment time points, a significant PAE × morphine interaction was observed (F₁,₈₆ = 132.77, p < 0.0001). Following the development of morphine-induced allodynia in PAE mice, a single injection of MCC950 or vehicle was administered at day 23 post-CCI. In both sexes, post hoc comparisons revealed a significant difference between non-allodynic PAE + minor CCI + morphine + MCC950 mice and allodynic PAE + minor CCI + morphine + vehicle mice at 90 minutes post-MCC950 injection, with allodynia reversal persisting at 24 hours (day 24; #p < 0.01 for both time points; **Figure 2A, C**). These data confirm the critical role of NLRP3 inflammasome activation in sustaining morphine-prolonged allodynia in PAE mice, regardless of sex.

### 3.3 Morphine-induced prolonged allodynia in PAE mice is associated with elevated NLRP3 activity, as indicated by increased Caspase-1 levels and other downstream pro-inflammatory molecules near the injured nerve, which are reduced by MCC950 treatment

Following behavioral assessments (**Figure 2**), tissues were collected, and the injured sciatic nerves were analyzed. Key molecular comparisons included: morphine-treated control (non-allodynic) vs. PAE (allodynic) mice, vehicle-treated (non-allodynic) PAE mice vs. allodynic PAE mice, and MCC950-treated pain-reversed PAE mice vs. allodynic PAE mice. Notably, at day 24 post-CCI, despite all mice being exposed to minor nerve injury, most treatment groups were non-allodynic due to spontaneous reversal or MCC950 treatment—except for the PAE + morphine + vehicle group, which remained allodynic (**Figure 2**). Levels of Caspase-1 activity were measured at the injury site (**Figure 3A**). In both males and females, significantly increased caspase-1 activity was observed in the allodynic group (PAE + morphine + vehicle) compared to non-allodynic controls: sac + morphine + vehicle ($p < 0.0001$) and PAE + vehicle + vehicle (##p < 0.0001). MCC950 treatment significantly reduced caspase-1 activity in morphine-treated PAE mice that displayed allodynia reversal (*p < 0.0001) in both sexes. Between-sex comparisons revealed significantly higher caspase-1 activity in males compared to females during morphine-prolonged allodynia (#p < 0.0001). In males and females, significantly increased levels of IL-1β protein (**Figure 3B)** were observed in the allodynic group (PAE + morphine + vehicle) when compared to other non-allodynic control groups, sac + morphine + vehicle ($ p < 0.0009) and PAE + vehicle + vehicle (## p < 0.004). In both sexes, MCC950 treatment dramatically reduced IL-1β protein in morphine-treated PAE mice that displayed allodynia reversal (* p < 0.04). Sciatic nerve mRNA levels of IL-1β, NLRP3, and TNF-α were also evaluated **(Figure 3C-E**). In females, morphine-induced allodynia in PAE mice was associated with higher levels of IL-1β mRNA (**Figure 3C**) compared to the non-allodynic control groups ($p < 0.0001). MCC950 treatment reduced IL-1β mRNA in morphine-treated PAE females (*p = 0.01). In females, there was a significant increase in TNF-α mRNA in allodynic nerve-injured morphine-treated PAE mice compared to vehicle-treated PAE mice (## p = 0.04). Similarly, in males, allodynic nerve-injured morphine-treated PAE mice displayed greater TNF-α mRNA levels compared to sac morphine-treated mice (p = 0.02) (**Figure 3D**). Significant sex differences in TNF-α mRNA levels were observed, with females displaying greater fold increases compared to males in nerve-injured morphine-treated PAE mice (# p = 0.04). Interestingly, MCC950 treatment reduced TNF-α mRNA in morphine-treated male PAE mice that displayed allodynia reversal (*p = 0.03). In males, morphine-induced allodynia was also associated with higher **NLRP3** mRNA levels (**Figure 3E**) compared to non-allodynic controls (p < 0.05). NLRP3 mRNA levels were significantly higher in male PAE mice compared to females during morphine-mediated allodynia (#p < 0.0001). MCC950 treatment significantly decreased NLRP3 mRNA levels in males compared to morphine-treated PAE mice with persistent allodynia (*p = 0.007). No NLRP3 mRNA changes were observed in females across treatment groups.

### 3.4 Morphine-induced prolonged allodynia in PAE mice is associated with upregulated NLRP3-related proinflammatory molecules in the spinal cord, which are reduced by MCC950 treatment

Ipsilateral lumbar spinal cord dorsal horn tissues were evaluated for molecular changes associated with TLR4–NLRP3 proinflammatory signaling. Caspase-1 activity and IL-1β protein levels were assessed to evaluate inflammasome activation in the spinal cord of both male and female mice. In males (**Figure 4B**), Caspase-1 activity was elevated in PAE + morphine + vehicle mice compared to sac + morphine + vehicle controls ($p = 0.004) and was reduced following MCC950 treatment (*p = 0.04). In females (**Figure 4A**), Caspase-1 activity was elevated in PAE + morphine + vehicle mice compared to PAE + vehicle + vehicle controls ($p < 0.0001; ##p < 0.0001). MCC950 treatment significantly reduced spinal caspase-1 activity in morphine-treated PAE mice (*p = 0.002). For IL-1β protein (**Figure 4C**), regardless of sex, morphine treatment significantly increased IL-1β levels in PAE mice compared to both sac + morphine + vehicle ($p < 0.005) and PAE + vehicle + vehicle (##p < 0.003) controls. MCC950 treatment significantly reduced spinal IL-1β protein levels in morphine-treated PAE mice (*p < 0.05). Higher IL-1β protein levels were observed in female PAE mice compared to males during morphine-mediated allodynia (#p = 0.0001). A similar pattern was observed at the transcript level. IL-1β mRNA (**Figure 5A**) was significantly increased in both male and female PAE + morphine + vehicle mice during morphine-prolonged allodynia, compared to sac + morphine + vehicle controls ($p < 0.0008) and PAE + vehicle + vehicle (##p < 0.03). In both sexes, MCC950-mediated allodynia reversal was associated with downregulation of spinal IL-1β mRNA (*p < 0.004). Morphine-prolonged allodynia also resulted in significant increases in TNF-α mRNA levels (**Figure 5B**) in both males and females compared to sac + morphine + vehicle controls ($p < 0.04). In females, spinal TNF-α mRNA was significantly higher in PAE + morphine + vehicle mice compared to PAE + vehicle + vehicle (##p = 0.006). In both sexes, MCC950-mediated allodynia reversal was associated with a decrease in TNF-α mRNA levels (*p < 0.03). In males, NLRP3 mRNA levels (**Figure 5C**) were significantly increased in the PAE + morphine + vehicle group compared to non-allodynic controls (sac + morphine + vehicle, $p = 0.02; PAE + vehicle + vehicle, ##p = 0.05). In females, NLRP3 mRNA was also elevated in allodynic PAE + morphine + vehicle mice compared to PAE + vehicle + vehicle controls (##p = 0.004). No significant changes in NLRP3 mRNA were observed following MCC950 treatment in either sex. Finally, morphine treatment significantly increased IL-18 mRNA levels (**Figure 5D**) in the spinal cords of both male and female PAE mice compared to sac + morphine + vehicle controls ($p < 0.03). In females, IL-18 mRNA was also significantly elevated compared to PAE + vehicle + vehicle mice (##p = 0.0002). MCC950 treatment significantly reduced IL-18 mRNA levels in both sexes, coinciding with behavioral reversal of allodynia (*p < 0.04).

**Figure 4.**
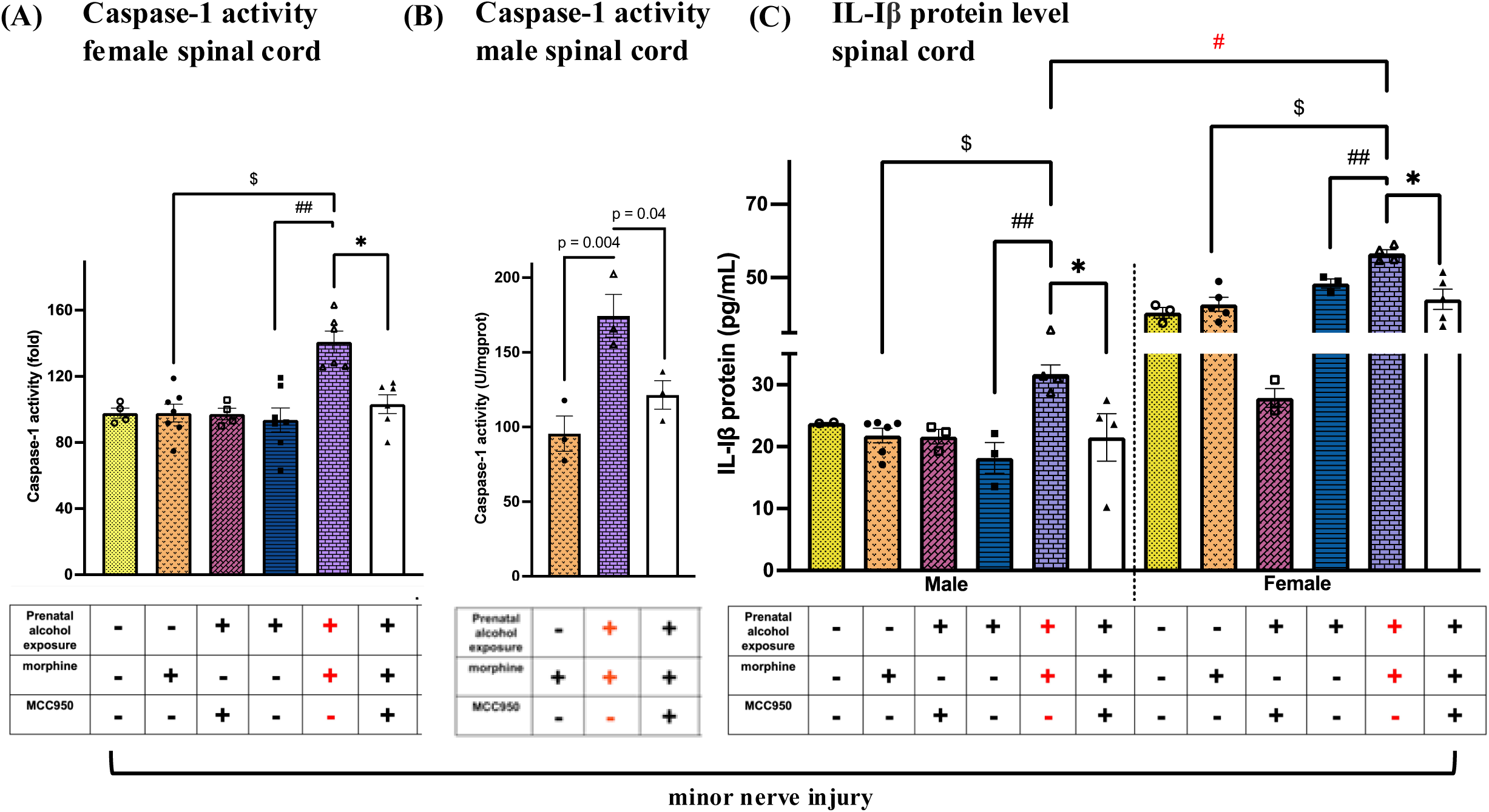
Effects of PAE and morphine immune interactions, and MCC950 treatment on Caspase-1 and IL-1β protein in the spinal cord from minor nerve-injured mice. Spinal cords were collected from behaviorally verified mice, as represented in Figure 2. Prenatal exposure (+), morphine (+), and MCC950 (-) in purple denote the allodynic group; all other groups are non-allodynic or allodynia-reversed at this post-CCI time point. **(A)** Despite all female mice being exposed to minor nerve injury, morphine treatment significantly increased spinal Caspase-1 activity in PAE compared to sac mice ($ p < 0.0001, PAE vs. sac, morphine-treated mice) and compared to PAE mice without morphine treatment (## p < 0.0001, PAE morphine vs. vehicle). MCC950 treatment reduced spinal Caspase-1 activity in morphine-treated PAE female mice that displayed allodynia reversa**l** (*p = 0.002). **(B)** In males, Caspase-1 activity was elevated between allodynic morphine-treated PAE mice and morphine-treated sac mice (p = 0.004, unpaired *t*-test). MCC950 treatment reduced spinal Caspase-1 activity in morphine-treated PAE male mice that displayed allodynia reversa**l** (p = 0.04, unpaired *t*-test). **(C)** Despite all mice being exposed to minor nerve injury, morphine treatment significantly increased IL-1β protein levels in PAE female and male mice compared to sac mice ($ p < 0.005, PAE vs. Sac, morphine-treated mice) and PAE mice without morphine treatment (## p < 0.003, PAE morphine vs. PAE vehicle). MCC950 treatment reduced spinal IL-1β protein in morphine-treated PAE mice that displayed allodynia reversa**l** (*p < 0.05). We observed significant sex differences in IL-1β protein, with females exhibiting higher levels compared to males in allodynic minor nerve-injured morphine-treated PAE mice (#p = 0.0001).

**Figure 5.**
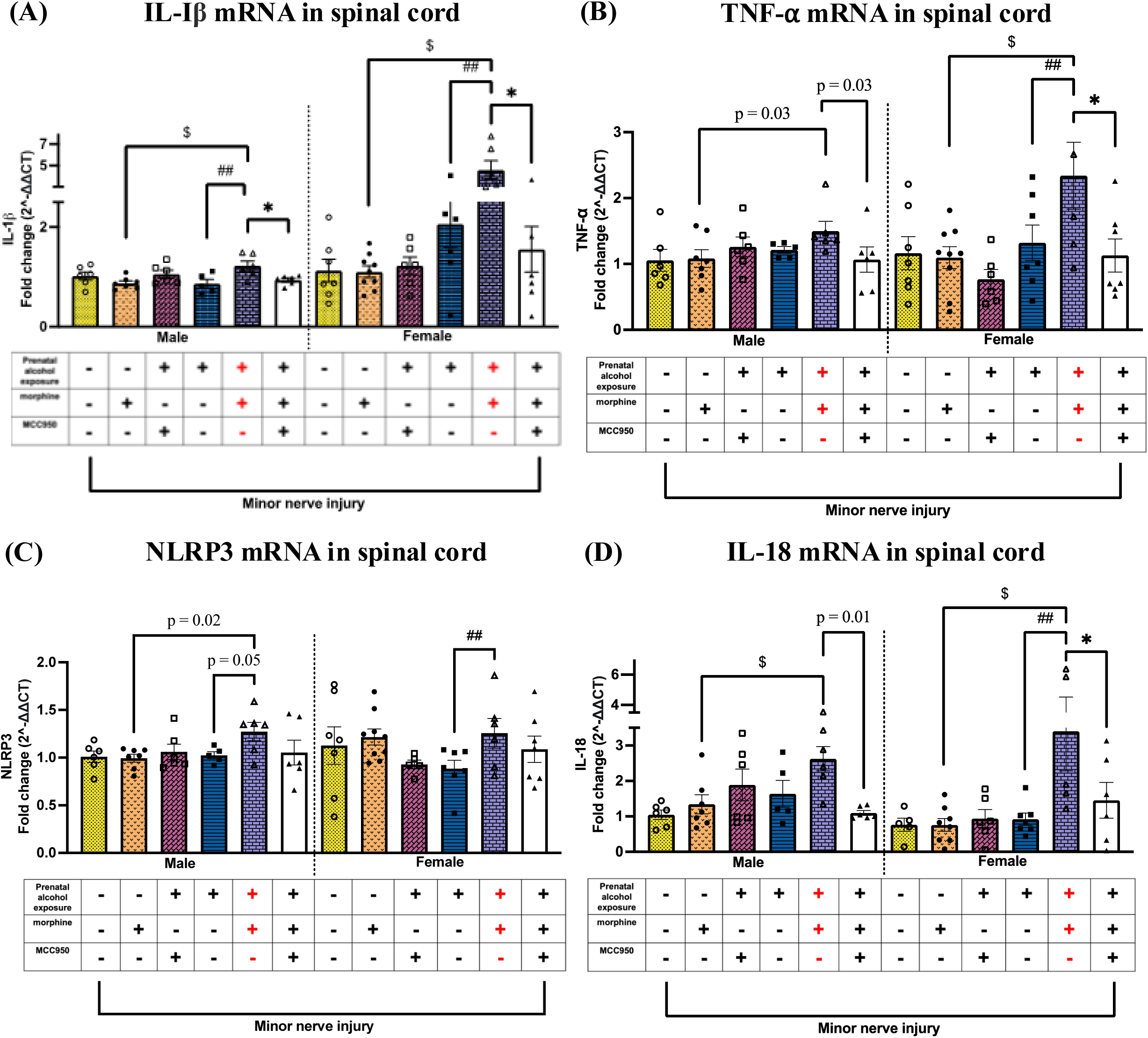
Effects of PAE, morphine interactions, and MCC950 treatment on IL-1β, NLRP3, and TNF-α mRNA levels in the spinal cord from minor-nerve injured mice. Spinal cords were collected from behaviorally verified mice, as represented in Figure 2. Prenatal exposure (+), morphine (+), and MCC950 (−) in purple denote the allodynic group; all other groups are non-allodynic or allodynia-reversed at this post-CCI time point. **(A)** Despite all mice being exposed to minor nerve injury, morphine treatment significantly increased IL-1β mRNA in both sexes in PAE mice compared to sac mice ($p < 0.0008, PAE vs. sac, morphine-treated mice) and PAE mice without morphine treatment (##p < 0.03, PAE, morphine vs. vehicle). Regardless of sex, MCC950 treatment reduced IL-1β mRNA in morphine-treated PAE mice that displayed allodynia reversal (*p < 0.004). **(B)** In males, morphine-prolonged allodynia coincided with a significant increase in TNF-α mRNA in PAE mice compared to sac mice (p = 0.03, unpaired t-test, PAE vs. sac, morphine-treated mice). In females, morphine-prolonged allodynia coincided with a significant increase in TNF-α mRNA in PAE mice compared to sac mice ($p = 0.0005, PAE vs. sac, morphine-treated mice) and PAE mice without morphine treatment (##p = 0.006, PAE, morphine vs. vehicle). Regardless of sex, MCC950 treatment significantly reduced TNF-α mRNA levels in morphine-treated PAE mice that displayed allodynia reversal (female: *p = 0.009; male: p = 0.03, unpaired t-test). **(C)** In males, morphine-prolonged allodynia coincided with a significant increase in NLRP3 mRNA in PAE mice compared to sac mice (p = 0.02, unpaired t-test, PAE vs. sac, morphine-treated mice) and PAE mice without morphine treatment (p = 0.05, unpaired t-test, morphine vs. vehicle). In females, regardless of minor nerve injury, morphine treatment significantly increased NLRP3 mRNA levels in morphine-treated PAE mice compared to vehicle-treated PAE mice (##p = 0.004). **(D)** In males and females, regardless of minor nerve injury, morphine treatment significantly increased IL-18 mRNA levels in PAE mice compared to morphine-treated sac male mice ($p < 0.03) and vehicle-treated PAE mice (female: ##p = 0.0002; male: p = 0.01, unpaired t-test). Regardless of sex, MCC950 treatment significantly reduced IL-18 mRNA levels in morphine-treated PAE mice that displayed allodynia reversal (*p = 0.03).

### 3.5 Morphine-induced prolonged allodynia in female PAE mice is associated with upregulated TLR4 and IκBα expression in the spinal cord, which is reduced by MCC950 treatment

We explored potential changes in TLR4 and IκBα mRNA levels in the spinal cord and sciatic nerve. Interestingly, while no changes were observed in male mice, spinal TLR4 mRNA levels (**Figure 6A**) were significantly increased in morphine-treated female PAE mice compared to non-allodynic sac + morphine + vehicle controls ($p = 0.01) and PAE + vehicle + vehicle mice (##p = 0.03). MCC950 treatment reduced spinal TLR4 mRNA levels in females (*p = 0.05). Similarly, spinal IκBα mRNA (**Figure 6B**) did not change in males regardless of PAE, morphine, or MCC950 treatment. However, in females, IκBα mRNA levels were significantly higher in allodynic PAE + morphine + vehicle mice compared to sac + morphine + vehicle controls ($p = 0.03). MCC950 treatment significantly reduced IκBα mRNA in female PAE mice (*p = 0.02). A comparison between sexes revealed that during morphine-induced allodynia, female PAE mice exhibited significantly greater fold increases in both TLR4 and IκBα mRNA compared to males (#p < 0.02). No significant differences in TLR4 or IκBα mRNA levels were observed across groups or sexes in the sciatic nerve (**Supplemental Figure 1B, C**). However, sciatic nerve TLR4 and IκBα mRNA fold changes were significantly higher in allodynic, minor nerve-injured, morphine-treated male PAE mice compared to their female counterparts (#p < 0.03$).

**Figure 6.**
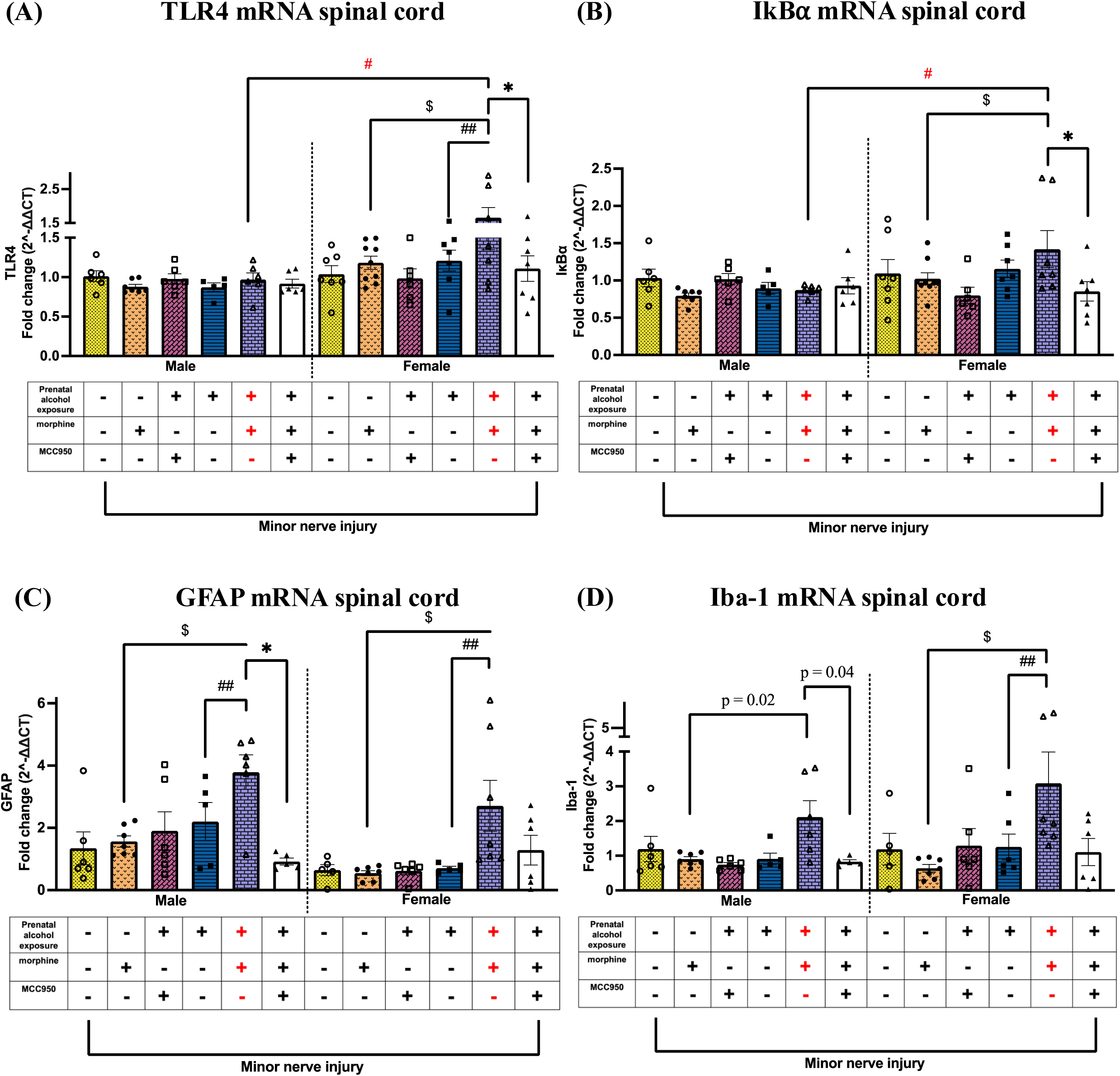
Effects of PAE, morphine and MCC950 treatment on TLR4 and IκBα mRNA levels in the spinal cord. **(A)** Morphine treatment increased TLR4 mRNA in the spinal cord only in allodynic PAE females compared to sac mice ($p = 0.01, PAE vs. sac, morphine-treated mice) as well as vehicle-treated PAE mice (##p = 0.03). MCC950 treatment reduced TLR4 mRNA in the spinal cord in female PAE mice that displayed allodynia reversal (*p = 0.05). No changes were observed in males. We observed significant sex differences in TLR4 mRNA, with females exhibiting higher levels compared to males in allodynic, minor nerve-injured, morphine-treated PAE mice (#p = 0.002). **(B)** Morphine treatment increased IκBα mRNA in the spinal cord only in allodynic PAE females compared to sac mice ($p = 0.03, PAE vs. sac, morphine-treated mice). MCC950 treatment reduced IκBα mRNA in the spinal cord in morphine-treated female PAE mice that displayed allodynia reversal (*p = 0.02). IκBα mRNA levels were significantly higher in allodynic female, minor nerve-injured, morphine-treated PAE mice than in their male counterparts (#p = 0.006). **(C)** Despite all mice being exposed to minor nerve injury, morphine treatment significantly increased GFAP mRNA in the spinal cord of allodynic PAE mice compared to sac mice regardless of sex ($p < 0.04, PAE vs. sac, morphine-treated mice) and PAE mice without morphine treatment (##p < 0.03, PAE, morphine vs. vehicle). In males, MCC950 treatment reduced GFAP mRNA in morphine-treated male PAE mice that displayed allodynia reversal (*p = 0.005). **(D)** Despite all mice being exposed to minor nerve injury, morphine treatment significantly increased Iba-1 mRNA in the spinal cord of allodynic PAE mice compared to sac mice regardless of sex (female: $p < 0.04; male: p = 0.02, unpaired t-test, PAE vs. sac, morphine-treated mice) and PAE mice without morphine treatment (##p < 0.03, PAE, morphine vs. vehicle). In males, MCC950 treatment reduced Iba-1 mRNA in morphine-treated male PAE mice that displayed allodynia reversal (p = 0.04, unpaired t-test).

### 3.6 Morphine-induced prolonged allodynia in PAE mice is associated with cell-specific increases in Iba-1 and GFAP expression in both sexes

Given the established roles of GFAP and Iba-1 as markers of astrocyte and microglial activation, respectively, we examined whether PAE and morphine interactions promote cell-specific neuroimmune changes associated with morphine-prolonged allodynia (**Figure 6C**). Regardless of sex, morphine-prolonged allodynia in PAE mice was associated with higher levels of GFAP mRNA compared to sac + morphine + vehicle mice ($p < 0.04) and PAE + vehicle + vehicle controls (##p < 0.03). In males, MCC950 treatment reduced GFAP mRNA, though not significantly (*p < 0.05). In females, morphine-prolonged allodynia significantly increased Iba-1 mRNA in PAE + morphine + vehicle mice compared to both sac + morphine + vehicle ($p < 0.05) and PAE + vehicle + vehicle controls (##p < 0.007; **Figure 6D**). In males, Iba-1 mRNA was significantly increased in morphine-treated PAE mice compared to sac + morphine + vehicle controls ($p = 0.02). In males, MCC950-mediated allodynia reversal was associated with reduced Iba-1 mRNA levels (*p = 0.04).

### 3.7 Upregulation of HMGB1 at the level of the injured nerve and spinal cord may contribute to morphine-induced prolonged allodynia in PAE mice, which is reduced following MCC950 treatment

Given evidence that spinal HMGB1 acts as an endogenous immune activator in morphine-induced hyperalgesia and allodynia under non-PAE conditions ^12,68^, we investigated whether PAE and morphine interactions further disrupt HMGB1 levels, potentially contributing to morphine-prolonged allodynia. Our data suggest that in both sexes, minor nerve injury led to a dramatic increase in sciatic nerve HMGB1 mRNA levels in morphine-treated PAE mice compared to morphine-treated sac controls ($p < 0.03) or vehicle-treated PAE mice (##p < 0.03; **Figure 7A**). MCC950-mediated allodynia reversal was associated with reduced HMGB1 mRNA levels in both sexes (*p < 0.005); significance in females was determined by a t-test, while the effect in males emerged from a post hoc ANOVA comparison, indicating a more robust difference. Additionally, sciatic HMGB1 upregulation revealed a sex difference: male PAE mice exhibited more pronounced HMGB1 elevation than female PAE mice in the presence of morphine-prolonged allodynia (#p = 0.0001). In addition to the increased HMGB1 expression at the peripheral nerve injury site, spinal cord HMGB1 mRNA levels were also significantly upregulated (**Figure 7B**) in both sexes compared to morphine-treated sac controls ($p < 0.0001) and vehicle-treated PAE mice (##p < 0.0003). Notably, MCC950-mediated allodynia reversal was associated with a significant decrease in spinal HMGB1 mRNA levels (*p < 0.0004).

**Figure 7.**
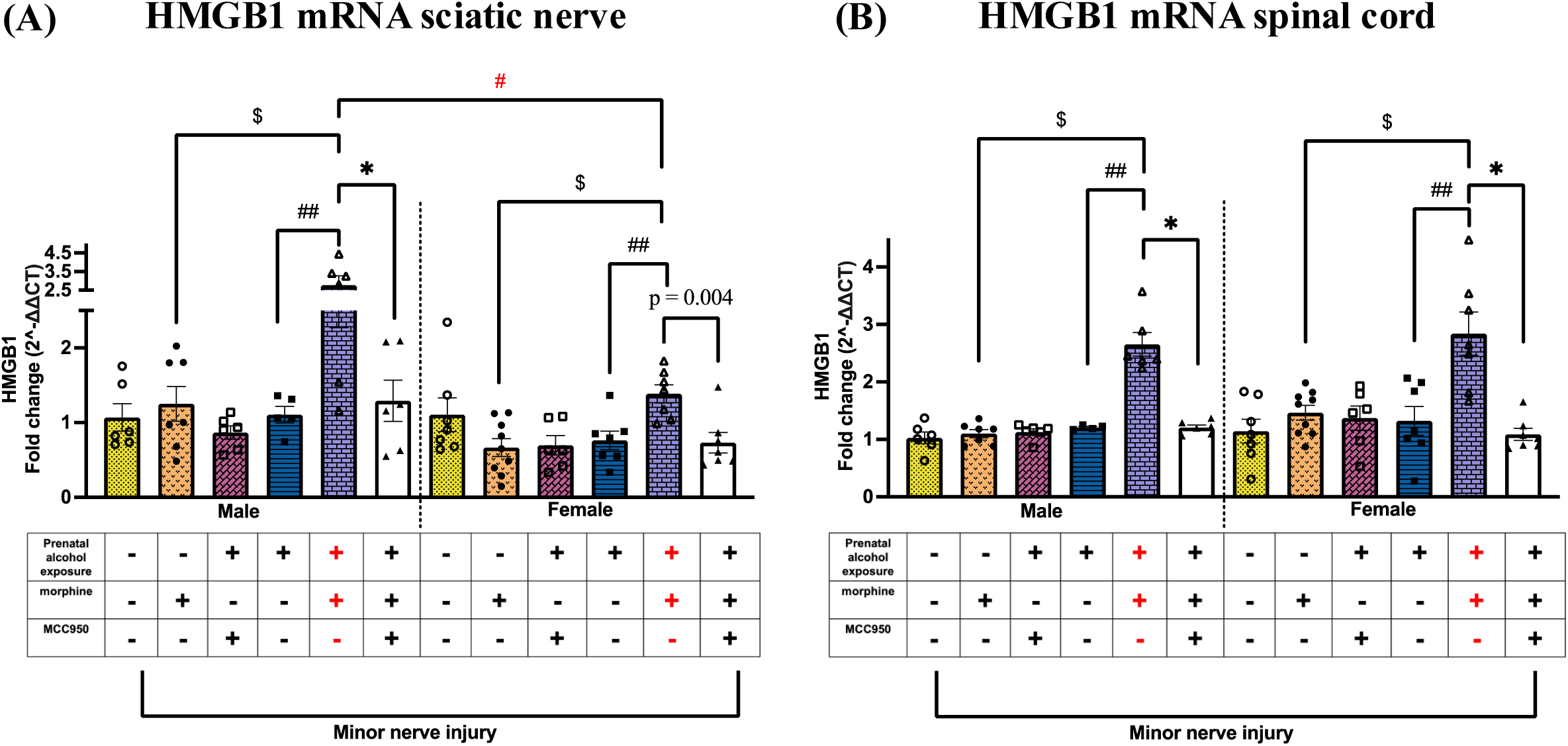
Effects of PAE and morphine and MCC950 treatment on HMGB1 mRNA levels in the injured nerve and spinal cord. **(A)** Despite all mice being exposed to minor nerve injury, and in both sexes, morphine treatment significantly increased HMGB1 mRNA in the sciatic nerve of allodynic PAE mice compared to sac mice ($p < 0.03, PAE vs. sac, morphine-treated mice) and PAE mice without morphine treatment (##p < 0.03, PAE, morphine vs. vehicle). MCC950 treatment reduced HMGB1 mRNA in morphine-treated male PAE mice that displayed allodynia reversal (male: *p < 0.002; female: p = 0.004, unpaired t-test). We observed that HMGB1 mRNA was significantly higher in allodynic male, minor nerve-injured, morphine-treated PAE mice compared to their female counterparts (#p = 0.001). **(B)** Independent of minor nerve injury, morphine treatment significantly increased spinal HMGB1 mRNA in both sexes in PAE morphine-treated mice compared to sac morphine-treated mice ($p < 0.0001, PAE vs. sac, morphine-treated mice) and PAE mice without morphine treatment (##p < 0.0003, PAE, morphine vs. vehicle). MCC950 treatment reduced HMGB1 mRNA in allodynia-reversed mice (*p < 0.0004).

### 3.8 Female PAE mice displayed increased spinal μ-opioid receptor mRNA levels during morphine-prolonged allodynia, which is decreased by MCC950 treatment

μ-Opioid receptor (MOR) mRNA was analyzed to assess potential modulatory effects of PAE, morphine, and MCC950 treatment. In both male and female PAE mice, morphine treatment significantly increased spinal μ-opioid receptor mRNA levels compared to morphine-treated sac controls ($p < 0.04; **Figure 8A**). In females, MCC950 treatment significantly reduced μ-opioid receptor mRNA levels (*p < 0.01). Notably, morphine-treated female PAE mice exhibited significantly higher fold changes in μ-opioid receptor mRNA expression in the spinal cord compared to their male counterparts (#p = 0.0007). No significant changes were observed in midbrain μ-opioid receptor mRNA levels across groups (**Figure 8B**).

**Figure 8.**
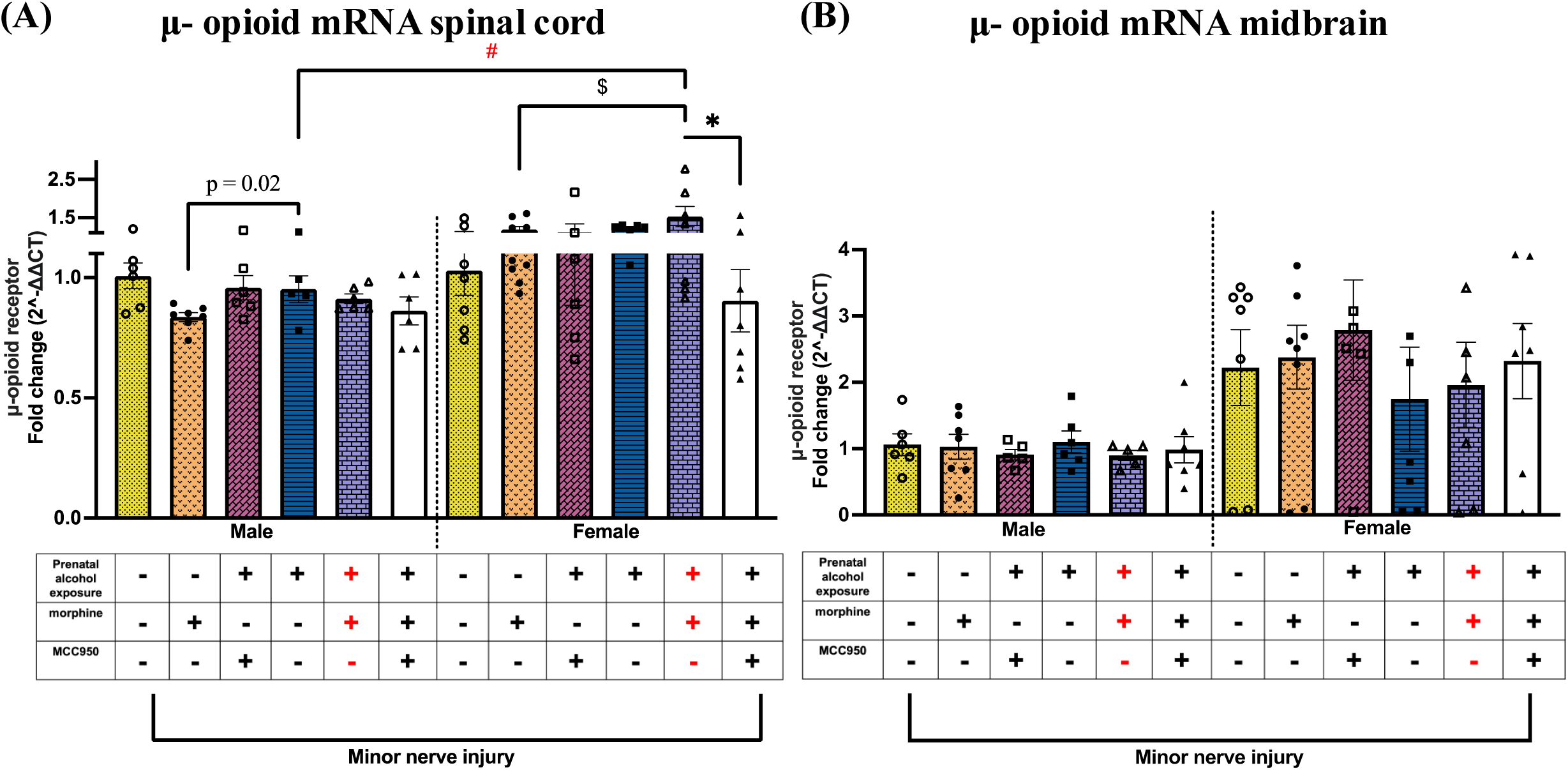
Effects of PAE and morphine immune interactions and MCC950 treatment on μ-opioid receptor mRNA in the spinal cord and midbrain in minor-nerve injured mice. **(A)** Regardless of sex, morphine-treated PAE mice exhibited significantly higher μ-opioid receptor mRNA levels compared to morphine-treated saccharin controls (female: $p = 0.03; male: p = 0.02, unpaired t-test). μ-Opioid receptor mRNA was significantly higher in allodynic female, minor nerve-injured, morphine-treated PAE mice compared to their male counterparts (#p = 0.0007). MCC950 treatment reduced HMGB1 mRNA in allodynia-reversed mice (*p < 0.01) **(B)** No change was observed in the midbrain.

### 3.9 Morphine-prolonged allodynia is associated with increased NLRP3 and TNF-α levels in different pain-relevant brain regions in females

To explore whether the interaction between PAE and morphine during nerve injury extends beyond the spinal cord and involves activation of TLR4 and NLRP3 signaling in pain-relevant brain regions, two key areas the midbrain and anterior cingulate cortex (ACC) were analyzed. In females, elevated NLRP3 mRNA levels were detected in the midbrain of PAE mice with morphine-induced allodynia compared to vehicle-treated PAE controls (##p = 0.02), while NLRP3 expression remained unchanged in males (**Figure 9A**). No significant changes were observed in IL-1β or TNF-α mRNA levels in the midbrain. In the ACC, allodynic PAE female mice exhibited significantly greater fold increases in TNF-α mRNA (**Figure 9B**) compared to non-allodynic, minor nerve-injured, vehicle-treated PAE controls (#p = 0.003). No MCC950-mediated changes were observed in any of these molecular markers in either the midbrain or ACC.

**Figure 9.**
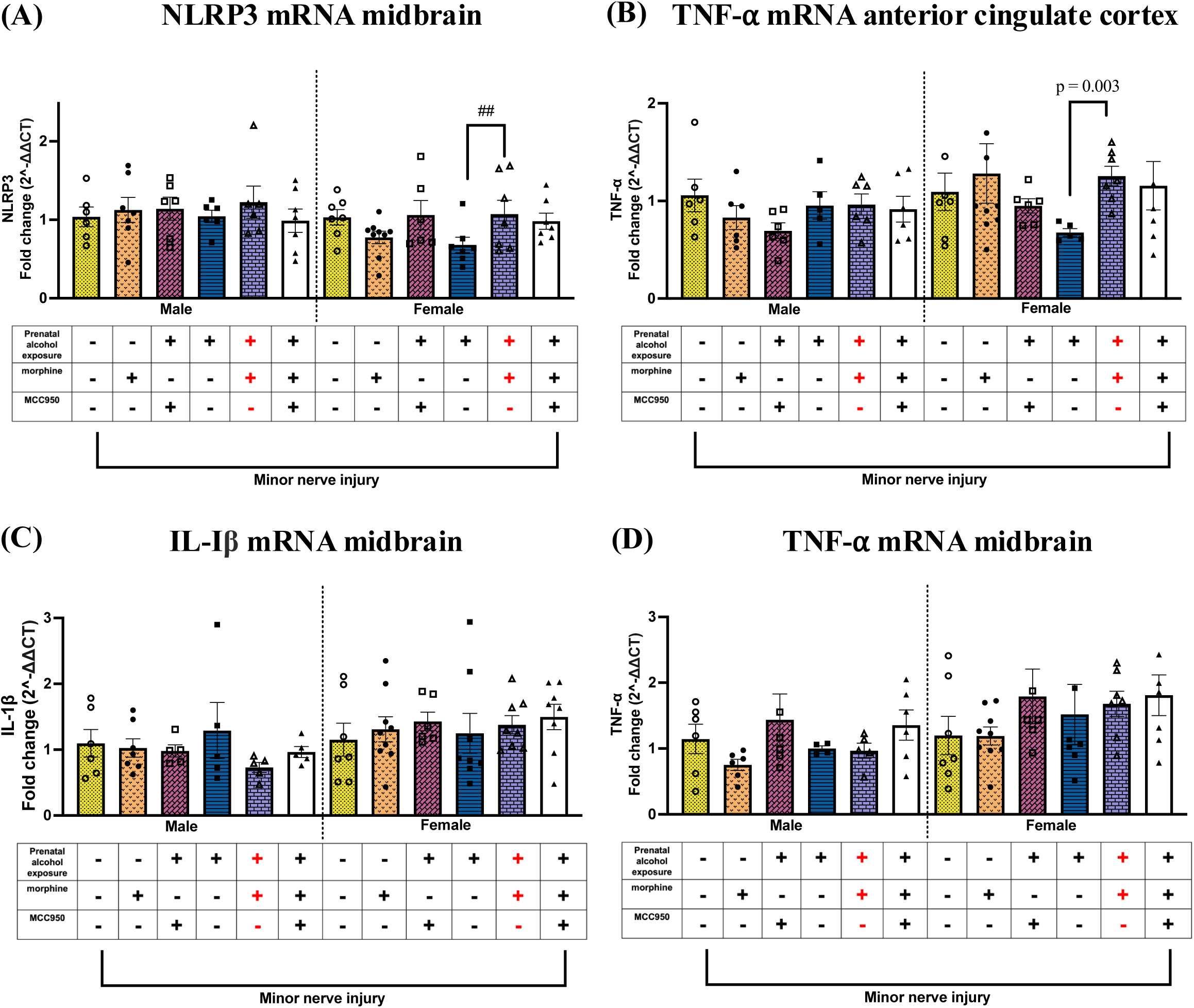
Effects of PAE and morphine and MCC950 on NLRP3 and TNF-α mRNA in the midbrain and anterior cingulate cortex. **(A)** Morphine treatment significantly increased NLRP3 mRNA in the midbrain of female PAE mice treated with morphine compared to PAE mice without morphine treatment (##p = 0.02). **(B)** Morphine treatment significantly increased TNF-α mRNA in the anterior cingulate cortex of female PAE mice treated with morphine compared to PAE mice without morphine treatment (p = 0.003, unpaired t-test). **(C & D)** No changes were observed in IL-1β or TNF-α mRNA in the midbrain regardless of sex.

**Figure 10.**
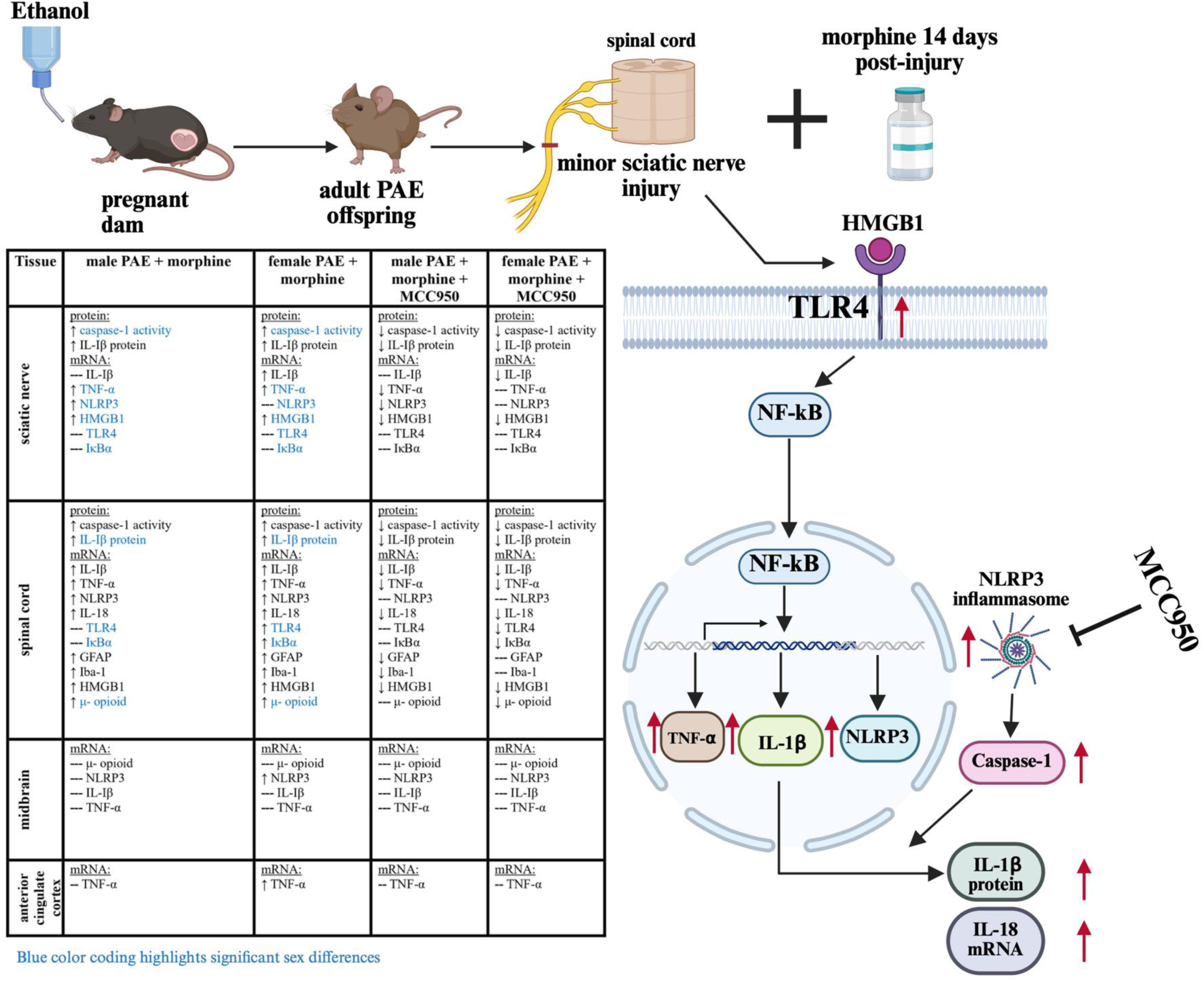
Schematic diagram of the experimental paradigm and summary of key findings of this study. This figure was created with Biorender.com

## 4. Discussion

Preclinical and clinical studies suggest neuroimmune dysregulation is a primary driver of FASD-related pathophysiology ^2,40,43,69^. TLR4 signaling modulates both immediate and long-term effects of PAE ^70^. Increased IL-1β has been reported due to alcohol exposure or PAE, following secondary immune activation ^41^. However, mechanistic insights confirming the necessary role of NLRP3 inflammasomes in PAE-induced CNS dysfunction are sparse. Our prior study demonstrated that systemic blockade of NLRP3, as well as blocking spinal IL-1β signaling ^41^, mitigates PAE-induced susceptibility to allodynia. Based on this strong scientific premise, we examined NLRP3 as a nexus of immune dysregulation consequent to PAE and opioid use in the context of nerve injury. While blocking NLRP3 activity with MCC950, which crosses the blood-brain barrier^71^, has been shown to reverse allodynia in female PAE mice ^44^, it remains unclear whether this prolonged allodynia results from peripheral and/or central immune signaling and how MCC950 exerts its beneficial effects. In line with our previous findings ^44^, this study generated novel behavioral and molecular data demonstrating that **(1)** PAE is a risk factor for morphine-induced allodynia in both male and female PAE mice, with a longer duration observed in males; **(2)** NLRP3 inflammasome activation is a key player in morphine and PAE neuroimmune interactions during nerve injury in both sexes; and **(3)** associated with upregulation of TLR4 and NLRP3-related factors in pain-relevant PNS and CNS regions, inhibiting NLRP3 activation reduces these critical proinflammatory factors, contributing to the reversal of allodynia.

### 4.1 PAE and morphine immune interaction during nerve injury drive NLRP3-dependent prolonged allodynia, regardless of sex

We find that morphine treatment significantly increases the chronicity of allodynia in minor nerve-injured PAE mice. Notably, morphine alone did not induce chronic allodynia in the absence of nerve injury, as evidenced by the lack of contralateral allodynia. Nonetheless, neither morphine nor PAE alone is sufficient to induce prolonged allodynia in the context of minor nerve injury. Instead, prolonged allodynia reflects immune sensitization arising from the interplay of these factors, unmasking PAE-induced susceptibility to morphine-mediated chronic pain outcomes. While our study examined the effects of morphine, similar effects may extend to other opioids, such as fentanyl and oxycodone, that activate TLR4 ^37,72^. Although unknown in the context of morphine-mediated allodynia, sex differences in allodynia ^73–75^ and PAE ^76–78^ have been reported in prior studies, as well as morphine^79^. Although we have observed a similar duration of minor nerve-induced allodynia in PAE mice without morphine treatment, these data are the first to suggest that the interaction between PAE and morphine results in a more extended duration of allodynia in males, compared to females.

### 4.2 PAE and morphine peripheral immune interaction involve heightened NLRP3 inflammasome activation at the site of injured nerve

Sciatic nerve injury causes localized nerve damage, which recruits heterogeneous immune cell populations, including macrophages, to the injury site ^11^. These macrophages release proinflammatory cytokines and mediators (e.g., HMGB1) that contribute to peripheral sensitization and enhancement of synaptic transmission in the dorsal horn of the spinal cord ^12^, facilitating the development and maintenance of central sensitization underlying chronic pain. Data presented in our study indicate that augmented peripheral immune activity with elevated Caspase-1 and other TLR4-related proinflammatory mediators engages in morphine-mediated allodynia under PAE conditions, consistent with primed activity of peripheral PAE macrophages ^44^. Moreover, MCC950 effectively reversed allodynia and reduced IL-1β and Caspase-1 levels, as well as mRNA levels of IL-1β and TNF-α. Although MCC950 acts at the level of “activation” of the NLRP3 inflammasome complex, which is downstream of NF-κB-mediated transcription of NLRP3 and IL-1β, TNF-α, these observations at 24 hours post-MCC950 treatment are likely due to suppression of cytokine signaling, disrupting this feed-forward loop of NF-κB activation^47,62,80,81^.

### 4.3. PAE and morphine immune interaction upregulated spinal glial activation and NLRP3– and TLR4-related factors following minimal nerve injury

The critical role of the spinal IL-1β signaling and activation of microglia and astrocytes is well documented during allodynia ^82,83^. Our data revealed increased levels of key mediators of allodynia-Caspase-1, IL-1β, NLRP3, and TNF-α and IL-18 ^28^, concurrent with spinal microglia and astrocyte activation in PAE mice with morphine-prolonged allodynia across both sexes ^43,84^. Reversal of allodynia following MCC950 treatment was associated with a reduction of these proinflammatory factors ^47,48^, as well as glial activation markers. Similar observations were made by prior studies showing NLRP3 inhibition attenuates GFAP mRNA and other neuroinflammatory factors during chronic pain states ^82,83^. MCC950 specifically blocks NLRP3 inflammasome ^57^, but not other inflammasomes ^57,85^ (such as NLRC4) that may contribute to Caspase-1 activity. Therefore, our data confirm the critical role of NLRP3-dependent Caspase-1 activity driving persistent allodynia in morphine-treated PAE mice, regardless of sex. Future studies examining the effects of NLRP3 inhibition at the spinal cord level could provide insights into whether ameliorating NLRP3 in both the PNS and/or CNS is necessary for allodynia reversal.

One critical aspect of this model to consider is that morphine is metabolized within a few hours to days^86^. Therefore, prolonged allodynia is likely to stem from morphine and PAE interaction that may enhance pain-promoting endogenous factors ^11^. Morphine has been shown to directly activate the immune cells via binding to MD2 ^87^ (an adaptor protein of TLR4) and may promote HMGB1 release, which may occur in both the peripheral immune or CNS glia, consistent with our prior findings on peripheral macrophages and the existing literature ^44,88,89^. At both the sciatic nerve and spinal cord levels, we have observed increased levels of HMGB1 in PAE mice, and the reversal of allodynia by MCC950 is associated with reduced HMGB1, suggesting that persistent allodynia in PAE mice may be attributed to ongoing TLR4 activation by HMGB1. These mRNA data are consistent with prior studies supporting increased levels of HMGB1 protein and confirming their critical contribution in the context of repeated morphine exposure resulting in chronic excitation of the TLR4-NF-kB axis ^35,36,90^. Further mechanistic approaches are needed to determine whether HMGB1 plays a necessary role in sustaining morphine-prolonged allodynia in PAE mice. In addition to morphine-mediated actions on immune cells, indirect mechanisms of immune modulation via interacting with the neuroendocrine system, can also be involved. Repeated exposure to morphine may blunt the hypothalamic-pituitary-adrenal (HPA) responsiveness and dysregulate immune function ^91,92^. Separately, PAE disrupts the HPA axis during adulthood, with hyperactive HPA in response to stress and a blunted response to cortisol associated with heightened proinflammatory factors ^93–96^ has been observed. Lastly, chronic pain can act as a stressor ^97–99^. Therefore, PAE and morphine interactions via the HPA axis are likely involved, which is a critical area to be explored in future studies.

### 4.4 Sex-specific responses during PAE and morphine-mediated immune interactions in the presence of nerve injury

During allodynia, variable spinal expression of NLRP3 and differential involvement of TLR4 has been reported ^26,43,100^. The critical role of the spinal Caspase-1 has been confirmed during morphine-mediated allodynia in non-PAE male rats ^36^. This study is the first to conduct a side-by-side comparison of TLR4-NLRP3-related proinflammatory molecules, revealing sexual dimorphism in PAE and morphine-mediated immune interactions. To summarize our molecular data common in both sexes, upregulation of Caspase-1 activity, HMGB1, and IL-1β was observed in both the periphery and the spinal cord, along with spinal glial activation in PAE mice with morphine prolonged allodynia. Moreover, MCC950 treatment reversed allodynia and dampened these factors in both sexes. Despite the critical role of NLRP3 activation regardless of sex, our molecular data revealed distinct inflammatory profiles in the peripheral versus CNS across the sexes. We found that PAE females display a more robust spinal immune sensitization than males along the TLR4-NF-κB-IL-1β pathway; elevated levels of TLR4, IκB-α, NLRP3, IL-1β, TNF-α, and IL-18 were observed. Interestingly, in the periphery, despite similar levels of IL-1β protein with males and upregulation of IL-1β and TNF-α mRNA levels, minimal changes in TLR4, IκB-α, and NLRP3 were observed in females. This disconnect may reflect the contribution of other NF-κB-independent mechanisms (such as MAPK/AP-1) that may contribute to the mRNA transcription of these cytokines, which are upstream molecules of NLRP3 activation. Also, differential post-transcriptional control of the stability of these mRNAs may occur ^101,102^. In contrast, in the periphery, male PAE mice showed more pronounced TLR4-related inflammatory factors ^42^ with comparatively greater fold increases of TLR4 and IκB-α mRNA levels than in females, and with significant increases in NLRP3 mRNA and IL-1β protein. Typically, IκB-α transcription occurs due to NF-κB activity, to provide negative feedback to its own activation ^103,104^. While no changes in IκB-α mRNA may infer a lack of robust ongoing NF-κB activity at the injury site in PAE females at this time point, in a chronic inflammation scenario, IKB is often blunted and dysregulated, contributing to sustained inflammation ^105,106^. Therefore, more targeted approaches, such as nuclear localization of NF-κB or IκB-α activity, may pinpoint the involvement of NF-κB or other critical transcription factors influencing transcriptional regulation of these factors. Moreover, TLR4-NF-κB-mediated transcription promotes NLRP3 actions and vice versa, however, transcriptional upregulation of all components of the inflammasome complex may not be required for NLRP3 actions, and could be temporally regulated during chronic inflammation and varies between sexes ^107–109^. Together, these data reflect potential differential effects of PAE and morphine on the TLR4-NF-κB activity across sexes and in the PNS vs CNS yet converging to downstream NLRP3 inflammasome-mediated actions in both sexes.

Notably, spinal μ-opioid receptor mRNA expression was upregulated in PAE females during prolonged allodynia, in a similar pattern of spinal TLR4 and IκB-α changes, consistent with prior reports indicating the involvement of TLR4–NF–κB signaling in regulating the transcription of μ-opioid receptors ^34,110^. Our initial characterization in pain-relevant brain regions ^24,111^ revealed that mRNA levels of most of these immune factors were unaltered. Interestingly, elevated mRNA levels of TNF-α in ACC, and NLRP3 in midbrain were detected in female PAE mice during morphine-prolonged allodynia. However, our dissection method captured PAG along with other adjacent midbrain nuclei and may not reflect subtle molecular alterations that may occur in males. Further investigation is required to gain a more comprehensive understanding of the role of NLRP3 within the pain-relevant brain regions.

Lastly, although we have conducted the molecular analysis at a time point when both male and female mice were stably allodynic, in light of our data suggesting a longer duration of allodynia in male PAE mice, greater magnitudes of spinal proinflammatory molecules were not evident in males. However, during morphine-prolonged allodynia, greater HMGB1 levels were observed in PAE males than in females at the sciatic nerve. HMGB1 may promote NLRP3 actions and allodynia via other TLR receptors as well as non-TLR pathways, leading to various other proinflammatory mediators ^112^, which may work synergistically in PAE males. These data may also indicate potential differential regulation of pain resolution mechanisms that may involve dysregulation of anti-inflammatory molecules ^113,114^.

## Limitations

Although our data validated protein levels of a few critical effector molecules, e.g., Caspase-1 and IL-1β, due to insufficient tissue material, we prioritized mRNA profiling of other factors, with a targeted approach to the signaling pathway of interest. We observed a similar pattern in Caspase-1 and IL-1β protein levels. Our molecular detection of IL-1β protein could detect both pro– and mature IL-1β levels. Lastly, our data suggest involvement of IL-1β and IL-18, however, the necessary role of IL-1β and/or IL-18 in morphine prolonged allodynia in PAE mice is yet to be established. Lastly, this study provides a glimpse of spinal astrocytes and microglial activation in this context; the relative contributions of different spinal glial cell types in PAE-induced inflammatory milieu are still speculative and require cell-type-specific assessments in the future.

## Conclusions

We conclude that PAE poses a risk factor for peripheral and central immune sensitization from later-life exposure to morphine as a chronic pain therapeutic. NLRP3 inflammasome activation is a central driver of worsened allodynia outcome in PAE conditions. Targeting NLRP3 may be beneficial in mitigating neuroimmune dysfunction and adverse effects of morphine, as well as neurobehavioral deficits associated with FASD.

**Supplemental Figure 1.**
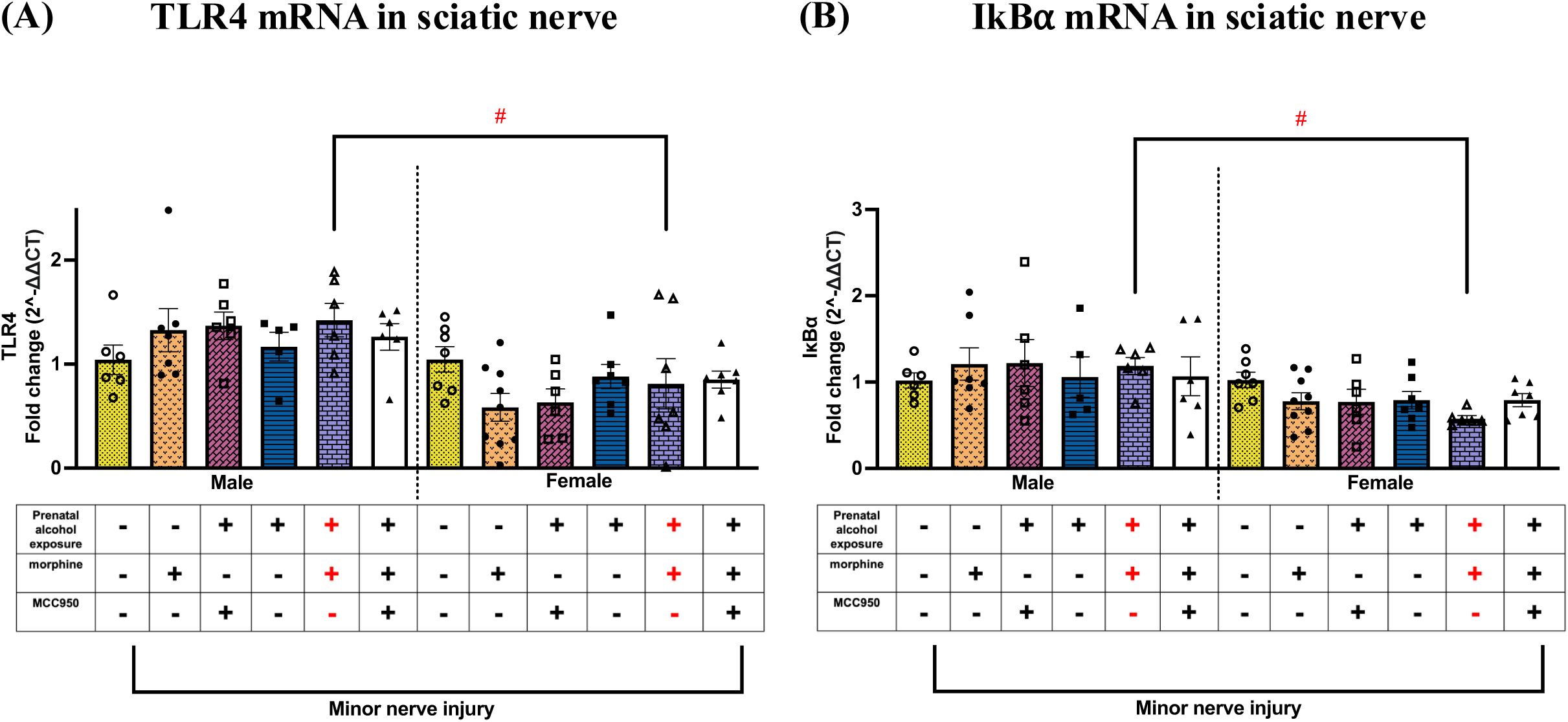
Effects of PAE and morphine and MCC950 Treatment on spinal protein IL-1β, TLR4, and IκBα mRNA levels in the sciatic nerve. **(A & B)** TLR4 and IκBα mRNA levels were significantly higher in allodynic male, minor nerve-injured, morphine-treated PAE mice compared to their female counterparts (#p < 0.03).

## Disclosures

### Funding

This study was funded by the National Institutes of Health/ NIAAA grants-R01 AA029694, R01 AA025967, and P50 AA022534.

## Conflicts of Interest

The authors declare that they have no competing interests in this study.

## Acknowledgments

The authors sincerely thank the UNM AIM core facility (NIH/P20GM121176) for providing access to the ELISA plate reader. We would like to acknowledge NIH URISE T34 GM145428 for access to Biorender. We would like to acknowledge Dr. Erin Milligan, Dr. David Linsenbardt & Dr. Sam McKenzie, UNM Neurosciences, for their useful advice in data analysis and interpretation.

## Data Availability Statement

All data generated and analyzed for the current report is included within the article.

## Authors’ Contributions

AAP and SN contributed to the conception and study design and prepared the manuscript. MS contributed to the behavioral and surgery training required for this manuscript. AAP and ANP performed behavioral assessments and data entry for hindpaw responses and morphine and MCC950 injections. AAP, SN, JC, and AKF collected tissue for molecular analysis. AKF aided with RNA standardization and reviewed data analysis. AAP performed molecular analysis of the sciatic nerve and spinal cord. AAP and AJ performed molecular analysis of brain regions. AAP performed statistical analysis for behavioral studies and molecular analysis. FV supervised the generation of moderate PAE-exposed offspring and aided in preparing the manuscript. DJ, MM, AAP, AKF, MS, and ANP performed the PAE paradigm. All authors contributed to the manuscript revision and read and approved the submitted version.

## Institutional Review Board Statement

The Institutional Animal Care and Use Committee (IACUC) of the University of New Mexico Health Sciences Center approved these studies (protocol # 21-201205-HSC, approval date 12/14/2021).

## Notes

### Competing Interest Statement

The authors have declared no competing interest.

### Summary of Updates

Corrected author sequence to match the original manuscript file and funder information.

